# The chromosomal passenger complex establishes chromosome biorientation via two parallel localization pathways

**DOI:** 10.1101/2020.09.10.288878

**Authors:** Theodor Marsoner, Poornima Yedavalli, Chiara Masnovo, Katrin Schmitzer, Christopher S. Campbell

## Abstract

Chromosome biorientation is established by the four-member chromosomal passenger complex (CPC) through phosphorylation of incorrect kinetochore-microtubule attachments. During chromosome alignment, the CPC localizes to the inner centromere, the inner kinetochore and spindle microtubules. Here we show that a small region of the CPC subunit INCENP/Sli15 is required to target the complex to all three of these locations in budding yeast. This region, the SAH, is essential for phosphorylation of outer kinetochore substrates, chromosome segregation, and viability. By restoring the CPC to each of these three locations individually, we found that inner centromere localization is sufficient to establish chromosome biorientation and viability independently of the other two targeting mechanisms. Remarkably, although neither the inner kinetochore nor microtubule binding activities was able to rescue viability individually, they were able to do so when combined. We have therefore identified two parallel pathways by which the CPC can promote chromosome biorientation and proper completion of mitosis.

## Introduction

To faithfully segregate their chromosomes, eukaryotic cells form a bipolar spindle with microtubules from opposite spindle poles attached to each of the two sister chromatids. Microtubules bind to the centromere region of chromosomes via a multi-protein complex called the kinetochore. During chromosome alignment prior to anaphase, many erroneous kinetochore-microtubule attachments are transiently formed. To maintain a stable genome from one generation to the next, it is imperative that cells detect and correct such erroneous attachments.

During mitosis, kinetochore-microtubule misattachments are detected and corrected by the evolutionally conserved chromosomal passenger complex (CPC). The CPC subunit Aurora B (Ipl1 in budding yeast) is a kinase that phosphorylates outer kinetochore substrates specifically on erroneously attached chromosomes (Welburn et al., 2010; Cimini et al., 2006; Tanaka et al., 2002; Lampson et al., 2004). This phosphorylation decreases the affinity of the kinetochore for the microtubules, allowing for detachment followed by reattachment in the correct orientation. How the CPC becomes proximal to the outer kinetochore and how it specifically targets misattachments are long-standing questions in the field (Lampson and Cheeseman, 2011; Krenn and Musacchio, 2015; Funabiki, 2019).

In addition to Aurora B/Ipl1, the CPC is composed of the subunits Survivin/Bir1, Borealin/Nbl1, and INCENP/Sli15. INCENP acts as a scaffold coordinating all of the CPC’s activities. The N-terminus binds to Survivin and Borealin, the C-terminus binds to and activates Aurora B, and the central region of the protein directly interacts with microtubules. The interactions with Survivin and Borealin facilitate targeting of the CPC to the inner centromere during chromosome alignment. This localization to the inner centromere is the predominant localization during early mitosis observed via microscopy, and was originally thought to be essential for the CPC’s role in chromosome biorientation. However, we previously found that a deletion mutant of Sli15 that eliminates the enrichment at the inner centromere is still able to undergo robust chromosome segregation in budding yeast (Campbell and Desai, 2013). Similar perturbations in human cells have recently been shown to maintain high levels of outer kinetochore phosphorylation by the CPC (Hadders et al., 2020). These results suggest that other mechanisms for targeting the CPC to the outer kinetochore exist. Indeed, recent evidence suggests that CPC-binding activities at the inner kinetochore can also contribute to outer kinetochore phosphorylation (Bonner et al., 2019). In yeast, inner kinetochore targeting occurs via a direct interaction between the central region of Sli15 and the COMA complex member Ctf19 (Fischböck-Halwachs et al., 2019; García-Rodríguez et al., 2019). Whether inner kinetochore binding of the CPC is sufficient to promote accurate chromosome segregation had not been previously investigated.

Microtubule binding by the CPC has also been shown to play a role in chromosome segregation. INCENP homologs contain a region directly upstream of the Aurora B-binding IN box that is predicted to form a single alpha helix (SAH) (van der Horst and Lens, 2013; Peckham and Knight, 2009). This region in chicken INCENP has been shown to be a single alpha helix that binds directly to microtubules *in vitro* (Samejima et al., 2015). In human cell culture, the SAH region contributes to centromere localization, kinetochore phosphorylation, and mitotic checkpoint activation (Wheelock et al., 2017; Vader et al., 2007).

In the budding yeast *S. cerevisiae*, the predicted SAH is part of a larger microtubule binding (MTB) domain in the central region of Sli15 (Kang et al., 2001). In addition to the SAH, the MTB contains a phospho-regulated (PR) region whose microtubule binding activity is suppressed by Cdk1 phosphorylation. At anaphase onset, the PR region is dephosphorylated by Cdc14 and the CPC relocalizes from the inner centromere to the microtubules of the mitotic spindle (Pereira, 2003). We have previously shown that the MTB is essential for CPC function in budding yeast, even though spindle localization occurs after chromosome biorientation, the only known essential function of the CPC in budding yeast, has been completed (Fink et al., 2017). Whether the essential function of this region is in microtubule binding, inner kinetochore localization, or some additional function is currently unknown.

In this study, we aimed to determine the function of the Sli15 SAH region in CPC-directed chromosome biorientation. We find that SAH mutants have reduced outer kinetochore phosphorylation, increased rates of chromosome missegregation, and are completely inviable. Surprisingly, mutations in the SAH region not only impair microtubule binding, but eliminate centromere and inner kinetochore CPC localization as well. By restoring targeting of the CPC to microtubules, the centromere and the inner kinetochore, we determined the contributions of each of these binding activities to CPC function. We found that inner centromere binding of the CPC was sufficient to partially rescue viability, whereas inner kinetochore binding was not. Intriguingly, combining microtubule binding with inner kinetochore binding resulted in rescue of CPC function, whereas the combination of microtubule binding and inner centromere binding had a strong synthetic negative affect. Our results therefore demonstrate the existence of two parallel pathways sufficient for targeting the CPC to the outer kinetochore to promote chromosome biorientation.

## Results

### The SAH region of Sli15 is essential for chromosome biorientation

A conserved feature of INCENP homologs is a predicted SAH region directly upstream of the IN box (van der Horst and Lens, 2013; Figure S1a). We previously demonstrated that large deletions of the *S. cerevisiae* INCENP homolog Sli15 that include the SAH are inviable (Fink et al., 2017). To determine whether the SAH region of Sli15 is essential for viability, we engineered Sli15 constructs that specifically disrupt the SAH either through deletion of this region or with point mutations. For the point mutants, twelve conserved basic residues in the SAH were mutated to glutamines in order to reduce the strong net positive charge of this region without disrupting the predicted helical fold (van der Horst et al., 2015; Figure 1a). Expression of the mutant constructs was confirmed by Western Blot (Figure S1b). After depletion of endogenous Sli15, both the Sli15-12Q and Sli15-ΔSAH constructs failed to rescue viability, indicating that the SAH region and its positively charged amino acids are essential for CPC function (Figure 1a). Additionally, both mutant constructs are lethal when overexpressed (Figure S1c), suggesting they act in a dominant negative manner to impair CPC function. We next measured chromosome segregation rates of Sli15-12Q and Sli15-ΔSAH, as chromosome biorientation is the only known essential function of the CPC in budding yeast (Biggins et al., 1999). In the first cell division following Sli15 depletion, GFP-labeled chromosome IV missegregated 62% of the time. Expression of Sli15-12Q or Sli15-ΔSAH only decreased the missegregation rates to 29% and 23% respectively, which is consistent with the observed lethality (Figures 1b and c). We conclude that the SAH region of Sli15 is essential for viability and faithful chromosome segregation.

**Figure 1.**
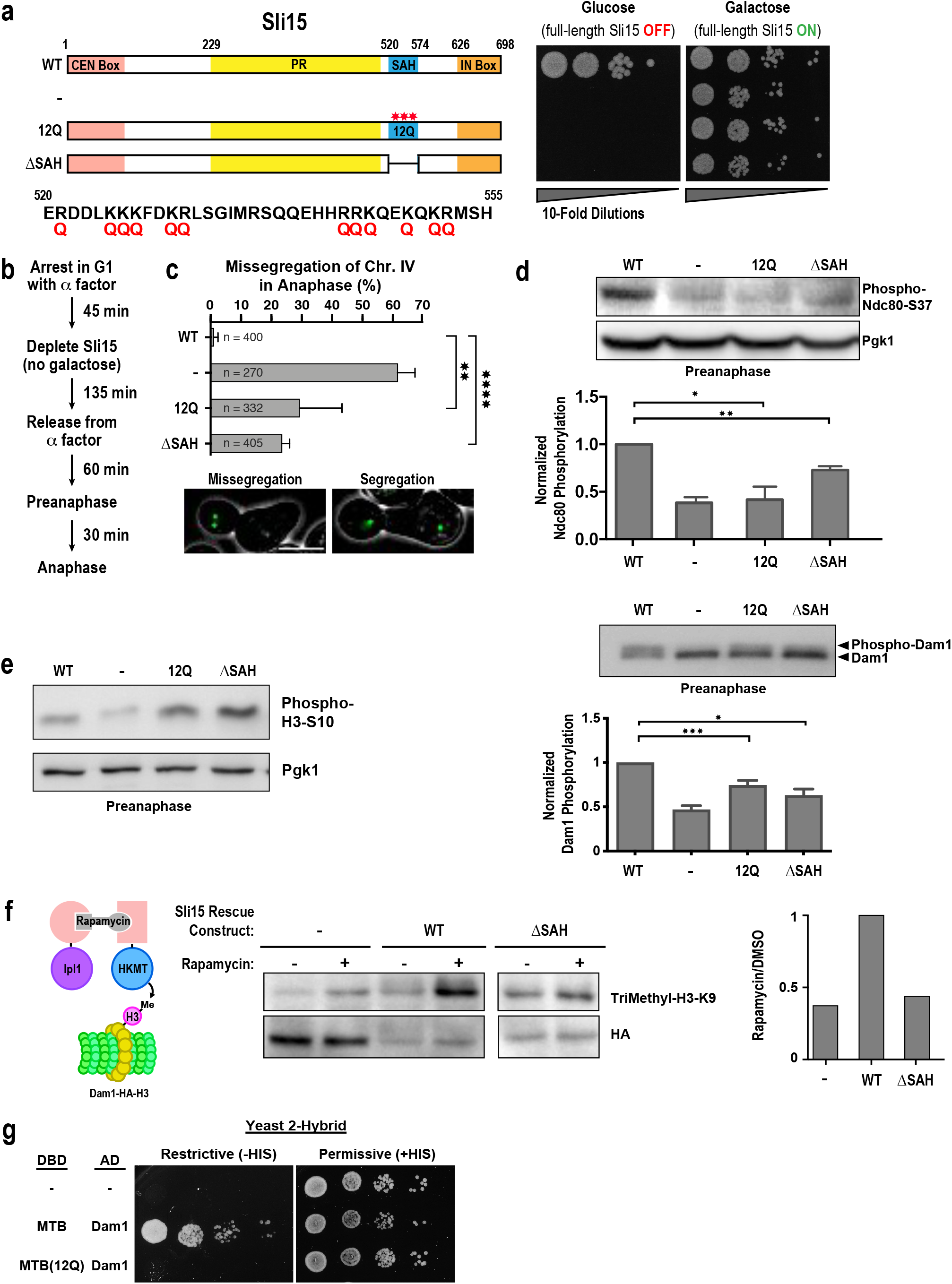
The SAH region of Sli15 is essential for chromosome biorientation and outer kinetochore phosphorylation. (**a**) 10-fold serial dilution analysis of cells expressing the indicated Sli15-SAH constructs with endogenous Sli15 under the control of a galactose inducible promoter. Schematic on the left shows the twelve glutamines (Q) mutations in the SAH region. (**b**) Protocol for Sli15 depletion in galactose deficient media and synchronization of cells with subsequent release from G1-arrest into anaphase. (**c**) Missegregation rates for GFP-labelled Chr. IV of indicated Sli15 constructs during the first anaphase after endogenous Sli15 depletion according to the protocol in **b**. Representative images show a missegregation event (left) and a correct segregation event (right). Scale bar: 5 μm. Mean and SD of at least three independent replicates are shown; (**d**) Immunoblotting analysis and quantification of Ndc80-S37 and Dam1 phosphorylation in preanaphase by the indicated Sli15 constructs. Cells were treated according to the protocol in **b**. Mean and SD of at least three independent replicates are shown. Pgk1 is shown as a loading control. (**e**) Immunoblot analysis of histone H3 serine 10 phosphorylation by the indicated Sli15 constructs in preanaphase cells. Phosphorylation levels of Sli15 mutants are not decreased (actually, slightly increased) with respect to wildtype Sli15. Pgk1 is shown as a loading control. (**f**) Immunoblot analysis and quantification for the proximity of indicated Sli15 constructs to Dam1 in preanaphase. Dimerization of Ipl1-FRB and HKMT-FKBP12 was induced by addition of rapamycin. Dam1 proximity was assessed via an antibody recognizing trimethylated signaling peptide H3-K9. HA-epitope present on the signaling peptide served as an internal loading control. (**g**) Yeast 2-Hybrid analysis of Sli15-MTB fused to DBD and Dam1 fused to AD of the Gal4 transcription factor. *: p < 0.05, **: p < 0.01, ***: p < 0.001, ****: p < 0.0001

### The SAH region of Sli15 is essential for outer kinetochore phosphorylation

The CPC facilitates chromosome biorientation by phosphorylating the outer kinetochore complexes Ndc80 and Dam1, which form direct kinetochore-microtubule attachments (Cheeseman et al., 2002; DeLuca et al., 2006; Cheeseman et al., 2006; Sarangapani et al., 2013). Ipl1-dependent kinetochore phosphorylation levels are high early in mitosis and decrease as cells become bioriented in metaphase (DeLuca et al., 2011; Keating et al., 2009). We defined the early “preanaphase” stage of mitosis as 60 minutes after G1 release. To detect Ipl1 phosphorylation of the kinetochore, we generated a phospho-specific antibody that recognizes phosphorylated serine 37 on Ndc80 and measured the signal by western blot (Figures 1d and S1d). In addition, we measured the phosphorylation-dependent mobility shift of Dam1. After synchronization and depletion of endogenous Sli15 in G1, we released the cells from the arrest and prepared preanaphase extracts (Figure 1b). Immunoblotting showed significantly decreased Ndc80 and Dam1 phosphorylation in cells expressing the Sli15-12Q and Sli15-ΔSAH mutants (Figure 1d). By contrast, phosphorylation of the conserved Aurora B/Ipl1 substrate Histone H3 on serine 10 was unchanged, demonstrating that global Ipl1 kinase activity is unaffected (Figure 1e) Together, these results demonstrate that the SAH region of Sli15 is specifically required for outer kinetochore phosphorylation by the CPC.

### The SAH region of Sli15 is essential for outer kinetochore proximity of the CPC

Decreased phosphorylation of CPC substrates at the outer kinetochore could result either from reduced proximity or diminished kinase activation specifically at that location. To detect kinetochore-proximal CPC independently of its activity, we used the M-track protein-protein proximity assay (Zuzuarregui et al., 2012; Brezovich et al., 2015). Ipl1 was bound to human histone lysine methyltransferase (HKMT) via a rapamycin inducible FRB-FKBP12 dimerization system. A short peptide from the HKMT substrate histone H3 was fused to Dam1 (Dam1-H3), which is irreversibly trimethylated if Ipl1-HKMT comes in close proximity (Figure 1f). Although the methylation site (K9) in this peptide is adjacent to an Ipl1 phosphorylation site (S10), methylation efficiency of this peptide was indistinguishable from an S10A mutant peptide (data not shown). CPC proximity to the outer kinetochore was determined by immunoblot using an antibody that specifically recognizes trimethylated histone H3. Upon rapamycin addition, a trimethylation signal was observed in the wild-type Sli15 strain in preanaphase. By contrast, Sli15-ΔSAH showed a relative trimethylation signal similar to that following depletion of Sli15, demonstrating that Sli15-ΔSAH is not proximal to the outer kinetochore (Figure 1f). The spatial resolution of the M-track assay was assessed by fusing the FRB tag to the centromeric protein Ndc10, which showed a strong trimethylation signal with the inner kinetochore protein Mif2-H3 but no rapamycin dependent increase in trimethlyation signal with Dam1. We therefore conclude that the M-track assay can distinguish between proximity to the inner vs. outer kinetochore (Figure S1e). The microtubule-binding (MTB) domain of Sli15 has previously been shown to interact with Dam1p in co-immunoprecipitation and yeast two hybrid assays (Kang et al., 2001). To test if this interaction requires a functional SAH region, we used the yeast two hybrid system with the DNA binding domain (DBD) of the transcription factor Gal4 fused to the MTB of Sli15 and the activation domain (AD) of Gal4 fused to Dam1. Wild-type MTB, but not MTB-12Q, was able to confer growth on media lacking histidine, demonstrating that a functional SAH is required for the interaction of the MTB with Dam1 (Figure 1g). Collectively, our results show that the SAH region, including the positively charged amino acids, is essential for viability, faithful chromosome segregation, outer kinetochore phosphorylation, and Dam1 proximity.

### The SAH region contributes to the spindle localization of Sli15

In vertebrates, the INCENP SAH region binds directly to microtubules and contributes to CPC association with the mitotic spindle (Samejima et al., 2015; van der Horst et al., 2015). To investigate the microtubule binding properties of the budding yeast SAH region directly *in vitro*, three MBP-tagged Sli15 microtubule binding domain constructs (MBP-MTB-WT, MBP-MTB-12Q and MBP-MTB-ΔSAH) were expressed and purified from *E. coli* and incubated with taxol-stabilized tubulin. Centrifugation and analysis of supernatant (S) and pellet (P) fractions via immunoblotting revealed a ~20% decrease of MBP-MTB-12Q and MBP-MTB-ΔSAH in the pellet fraction, indicating that the SAH region contributes to microtubule binding in vitro, although not as strongly as the PR region of the MTB (Figure 2a). To test whether the SAH region is necessary for the spindle association of the CPC in *S. cerevisiae*, Sli15-12Q and Sli15-ΔSAH were labeled with mNeonGreen and localization to the spindle was assessed both prior to, and shortly after anaphase onset. Both Sli15-12Q and Sli15-ΔSAH showed reduced localization to the preanaphase spindle compared to wild-type Sli15 (Figure 2b). Tub1 signal intensities were similar in all of the Sli15 constructs, indicating that the differences in intensity were not due to indirect effects of microtubule stability (Table S1). However, due to their close proximity, centromere or kinetochore binding could also contribute to the observed preanaphase localization signal. To eliminate any contribution to fluorescent signal from the centromeres or kinetochores, we measured the spindle localization of Sli15-12Q and Sli15-ΔSAH also in anaphase where the microtubules are more easily distinguished from centromeres and kinetochores. The mutant SAH constructs showed decreased localization to the spindle in anaphase, although this decrease was substantially more prominent in preanaphase (Figure 2b). We note that the contribution of the PR region of the MTB to spindle localization in anaphase is greatly enhanced by dephosphorylation of Cdk1 sites. This increased contribution by the PR region is also likely to occur in the MBP-MTB-12Q and MBP-MTB-ΔSAH constructs purified from bacteria, since *E. coli* lacks Cdk1 (Figure 2a). The *in vitro* microtubule binding assays are therefore comparable to the anaphase localization results. We conclude that the SAH region contributes only moderately to microtubule binding *in vitro* and in anaphase. The reduction of spindle association in preanaphase by Sli15-12Q and Sli15-ΔSAH could be due to an increased contribution of the SAH to microtubule binding relative to the PR region. Alternatively, the SAH could contribute to the enrichment of the CPC at other spindle/chromatin locations such as the centromeres or kinetochores.

**Figure 2.**
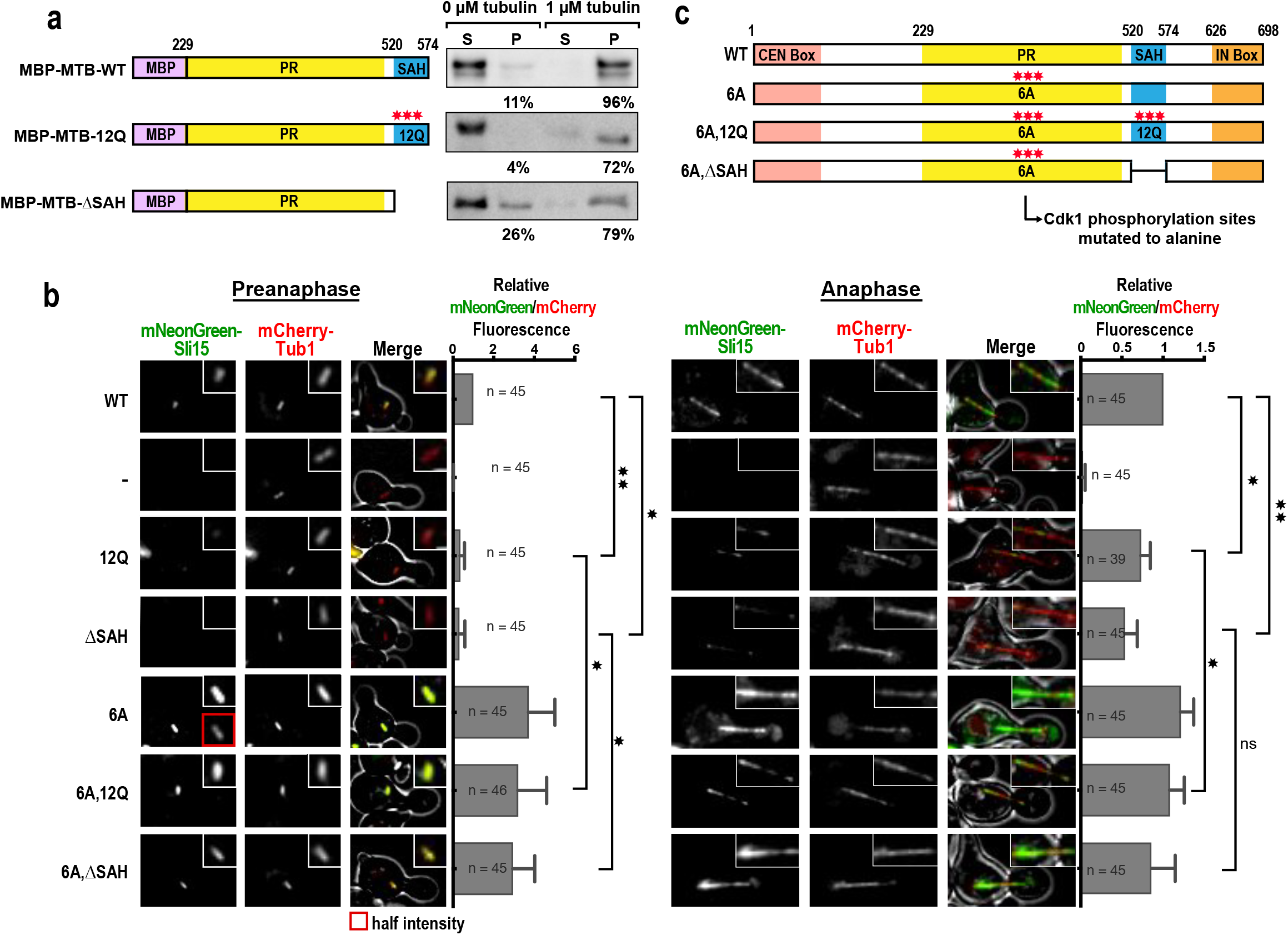
The SAH region of Sli15 contributes to microtubule binding and spindle association. (**a**) MT pelleting assay with indicated constructs expressed from *E. coli*. Constructs were incubated with 1 μM taxol-stabilized tubulin, centrifuged and MT pellets were analyzed using immunoblotting with an antibody to MBP. (**b**) Spindle localization of Sli15 constructs in preanaphase (left) and anaphase (right). Preanaphase spindles were defined as shorter than 2 μm and located entirely in the mother cell. Images of the same fluorophore were contrast adjusted identically. Note that the mNeonGreen signal of 6A in preanaphase was saturated, so an insert showing half the intensity was added for comparison. Spindle localization was quantified by fitting a Gaussian curve to a line perpendicular to the spindle center for both mNeonGreen and mCherry. Ratios of the integrated Gaussian fits from the mNeonGreen and mCherry signals were calculated. Mean and SD from three independent experiments are shown. For the values of individual fluorophores, see Table S1. Insert box preanaphase: 2.6 μm square; insert box anaphase: 6.25 μm in length. *: p < 0.05; **: p <0.01. (**c**) Schematic of Sli15 constructs containing six Cdk1 phosphorylation sites mutated to alanine in the PR region in combination with 12Q in the SAH and ΔSAH mutants.

### CPC spindle localization is sufficient for outer kinetochore phosphorylation but not chromosome segregation

If the essential function of the SAH is to localize the CPC to microtubules in preanaphase, then mutations that increase microtubule binding should compensate for the lack of a functional SAH region. To test this, six Cdk1 phosphorylation sites in the PR region were mutated to unphosphorylatable alanines (Sli15-6A). These mutations have been shown to prematurely target the CPC to mitotic spindle microtubules prior to anaphase. Importantly, the 6A mutant does not affect cell cycle timing, chromosome missegregation rates or viability (Pereira, 2003; Mirchenko and Uhlmann, 2010). We therefore combined the Sli15-6A mutations with Sli15-12Q or Sli15-ΔSAH to assess the function of microtubule binding independently of other possible functions of the SAH region (Figure 2c). Sli15-6A, Sli15-6A,12Q and Sli15-6A,ΔSAH all had stronger preanaphase spindle localization than wild-type Sli15 (Figure 2b). In anaphase, this difference was less pronounced, as the PR region is largely dephosphorylated at this stage. We conclude that the Sli15-6A mutant targets the CPC to the spindle microtubules even in the absence of the SAH region.

We next determined if microtubule binding is sufficient to rescue CPC function in the absence of the SAH region. Cells expressing Sli15-6A,12Q or Sli15-6A,ΔSAH double mutations that restore microtubule binding in the absence of the SAH were not viable following depletion of endogenous Sli15 (Figure 3a). In addition, the chromosome missegregation rates of the Sli15-6A,12Q and Sli15-6A,ΔSAH constructs, although showing strong preanaphase spindle localization, were similar to those of the Sli15-12Q and Sli15-ΔSAH constructs that show little-to-no spindle localization (Figure 3b). These results suggest that even though the SAH region contributes to spindle association in preanaphase, microtubule binding is not sufficient for the essential function of the SAH.

**Figure 3.**
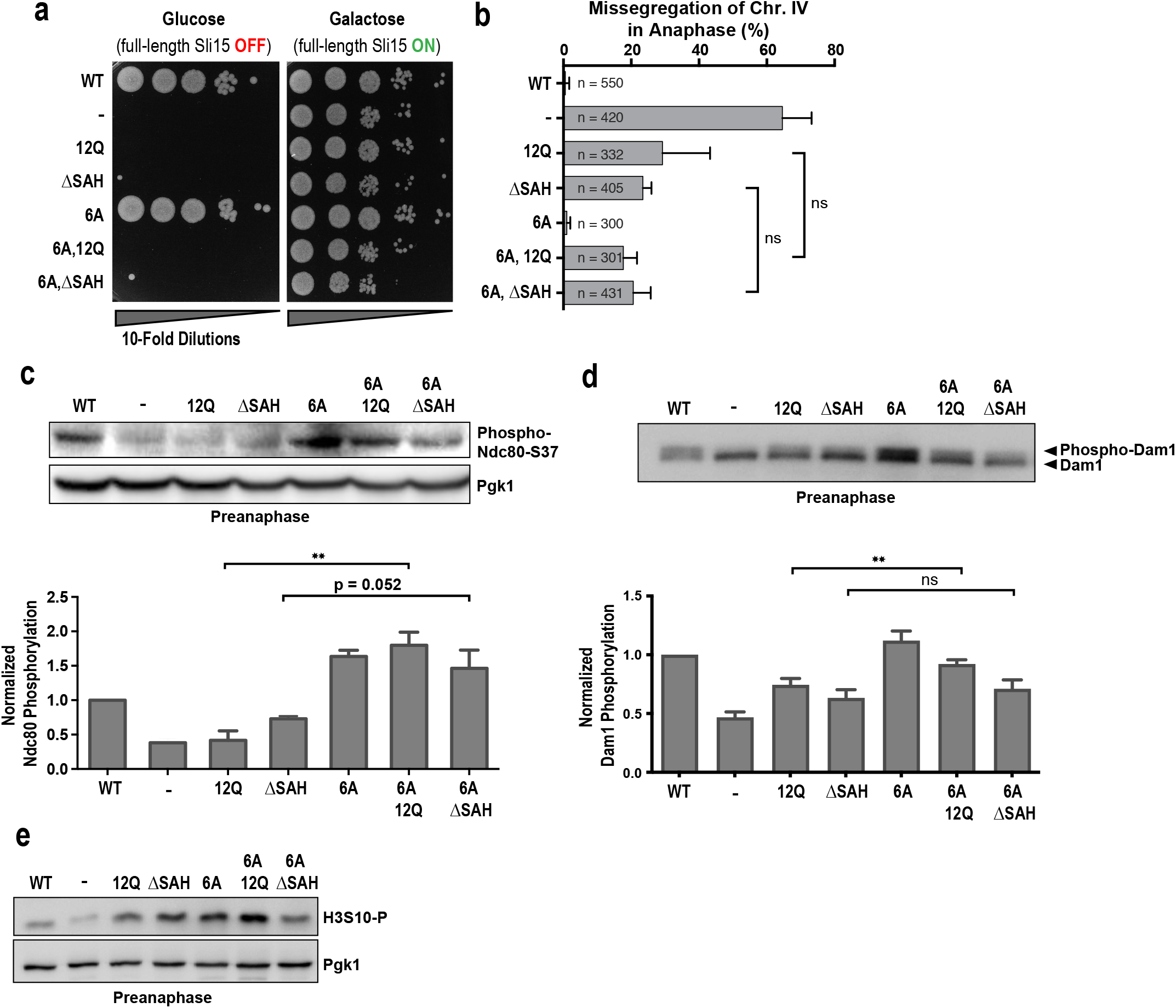
Microtubule binding by the CPC contributes to kinetochore phosphorylation but is not sufficient for biorientation. (**a**) 10-fold serial dilution analysis of indicated Sli15 construct expressing cells having endogenous Sli15 under the control of a galactose inducible promoter. (**b**) Missegregation rates for GFP-labelled Chr. IV of indicated Sli15 constructs during the first anaphase after endogenous Sli15 depletion as depicted in 1b. (**c**) Immunoblotting analysis and quantification of Ndc80-S37 in preanaphase by the indicated Sli15 constructs. Cells were treated as depicted in 1b and harvested in preanaphase. Mean and SD of at least three independent replicates are shown. Pgk1 is shown as a loading control. (**d**) Immunoblotting analysis and quantification of Dam1 phosphorylation in preanaphase by the indicated Sli15 constructs. Cells were treated as depicted in 1b and harvested in preanaphase. Mean and SD of at least three independent replicates are shown. Pgk1 is shown as a loading control. (**e**) Immunoblot analysis of histone H3 serine 10 phosphorylation by the indicated Sli15 constructs in preanaphase cells. Pgk1 is shown as a loading control. Data for WT, -, 12Q and ΔSAH in Figure 3 are reproduced from Figure 1. **: p < 0.01

We hypothesized that the inability of microtubule binding alone to promote chromosome biorientation results from a lack of outer kinetochore phosphorylation. Surprisingly, Ndc80 phosphorylation in preanaphase was even stronger in both Sli15-6A,12Q and Sli15-6A,ΔSAH than wild-type Sli15 (Figure 3c). In addition, Dam1 phosphorylation was fully rescued to wild-type levels by Sli15-6A,12Q (Figure 3d). The Sli15-6A, Sli15-6A,12Q and Sli15-6A,ΔSAH were all able to robustly phosphorylate Histone H3, indicating that global Ipl1 activity was not impaired in these mutants (Figure 3e). We conclude that CPC spindle localization contributes to outer kinetochore phosphorylation in preanaphase. Intriguingly, however, rescue of outer kinetochore phosphorylation by increased spindle association of the CPC is not sufficient to rescue error correction and viability.

### Microtubule-bound CPC phosphorylates kinetochores in a tension-independent manner

Current models for CPC-mediated error correction suggest that kinetochores respond to a lack of tension when misattached. When tension is applied by microtubule pulling forces at bioriented chromosomes, kinetochores are not phosphorylated and therefore remain stably attached to microtubules (reviewed in Lampson and Cheeseman, 2011). To determine how increased CPC spindle association leads to outer kinetochore phosphorylation without increasing the accuracy of chromosome segregation, we tested two hypotheses. First, excessive outer kinetochore phosphorylation could cause continuous disruption of kinetochore-microtubule attachments, including bioriented attachments that are under tension, leading to the observed chromosome missegregation and lethality. The unattached kinetochores created by excessive microtubule detachment would also result in a spindle assembly checkpoint-dependent arrest (Muñoz-Barrera and Monje-Casas, 2014). Even though the Ndc80 phosphorylation levels in Sli15-6A and Sli15-6A,ΔSAH cells were elevated, budding index measurements showed no accumulation of large-budded, metaphase-arrested cells (Figure 4a). Importantly, both mutants showed a strong metaphase arrest when treated with microtubule depolymerizing drugs, demonstrating that the spindle assembly checkpoint was still functional (Figure S2a). These results indicate that increased Ndc80 phosphorylation by Sli15-6A and Sli15-6A,ΔSAH does not lead to continuous kinetochore-microtubule detachment.

**Figure 4.**
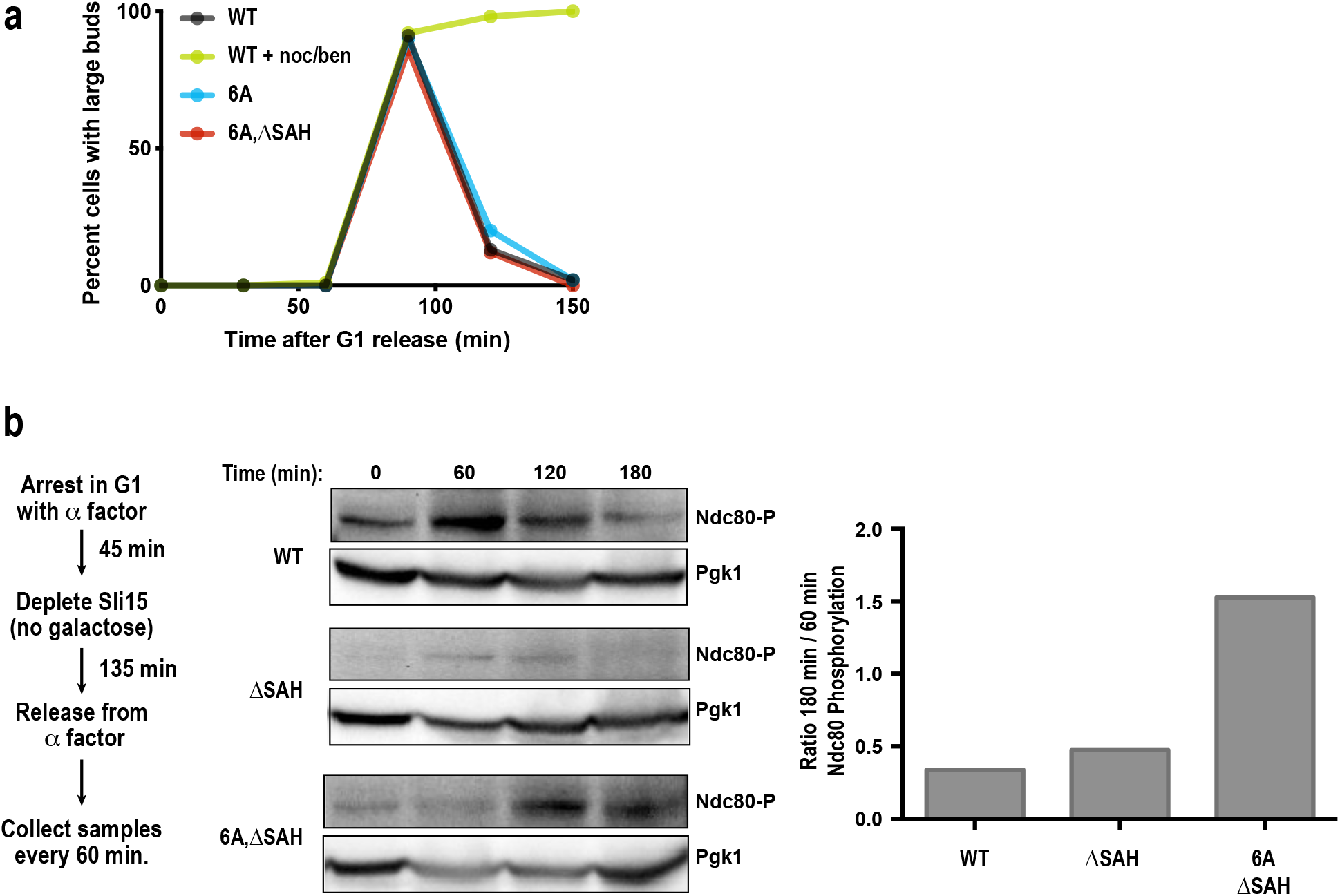
Microtubule binding by the CPC contributes to the phosphorylation of kinetochores in a tension-independent manner. (**a**) Budding index analysis of indicated Sli15 constructs. Cells were arrested in G1 and endogenous Sli15 was depleted. Fractions of big budded cells were determined every 30 min after release. Addition of nocodazole (5 μM) and benomyl (34 μM)) to WT Sli15 was used as a positive control for metaphase arrest. (**b**) Immunoblots and quantification of Ndc80-S37 phosphorylation over time. Cells containing the indicated Sli15 constructs were treated as indicated on the left. Pgk1 is shown as loading control. The ratio of Ndc80 phosphorylation between the 180 min and 60 min time points is plotted on the right. Quantification shows averages of two independent experiments.

Although no checkpoint activation was observed when expressed at endogenous levels, Sli15 mutants that increase microtubule binding (6A and ΔNT) exhibit a strong cell cycle arrest and lethality when overexpressed (Figure S2b). This finding indicates that CPC mutants with increased microtubule binding phosphorylate all kinetochores independently of the amount of tension and attachment state. Based on this observation, in the second hypothesis, we reasoned that when expressed at endogenous levels, the overall increase in phosphorylation levels seen for Sli15-6A,12Q and Sli15-6A,ΔSAH could result from a low level of phosphorylation at all kinetochores instead of a high level specifically at misattached kinetochores. This low level of phosphorylation would not be sufficient to detach microtubules from kinetochores. To test this theory, Ndc80 phosphorylation was monitored in cells over time from G1 to a Cdc20-depleted metaphase arrest (Figure 4b). Wild-type cells showed a strong accumulation of phosphorylation signal that peaked in preanaphase (60 min) and then decreased again by the 180-minute time point. A similar result was seen for cells with Sli15-ΔSAH, only with lower overall levels of phosphorylation. Strikingly, Ndc80 phosphorylation levels continued to increase in Sli15-6A,ΔSAH expressing cells at time points when chromosome biorientation should have already completed (Figure 4b). Indeed, phosphorylation of these mutants was even higher at the 180-minute time point than at the 60-minute time point. The aforementioned budding index measurements suggest that these differences are not attributable to changes in cell cycle timing (Figure 4a). We conclude that the spindle binding activity of the CPC contributes to tension-independent outer kinetochore phosphorylation and that this activity is not sufficient for substantial destabilization of kinetochore-microtubule attachments when expressed at endogenous levels.

### The SAH region of Sli15 is essential for inner centromere and inner kinetochore localization of the CPC

The above results show that although the SAH region contributes to microtubule binding of the CPC, this activity is not sufficient to compensate for the function of this region. We also found that localization to preanaphase spindles is almost completely absent in SAH mutants, suggesting that all CPC targeting activities are disrupted (Figure 2b). We therefore sought to determine if the SAH mutants disrupt other known targeting mechanisms of the CPC. To test whether the SAH contributes to inner centromere enrichment in budding yeast, cells were treated with a combination of nocodazole and benomyl to eliminate microtubules and promote the Sgo1-dependent accumulation of the CPC to the inner centromere (Verzijlbergen et al., 2014; Campbell and Desai, 2013). As previously observed, the localization of the CPC was enriched between the two kinetochore clusters upon drug treatment, with peak fluorescence increasing ~ 3-fold over the DMSO control (Figure 5a). This inner centromeric localization was undetectable in the Sli15-ΔSAH mutant cells. Nuf2 signal was largely unaffected by either Sli15 depletion or the Sli15-ΔSAH mutant, demonstrating that kinetochore formation still occurred normally.

**Figure 5.**
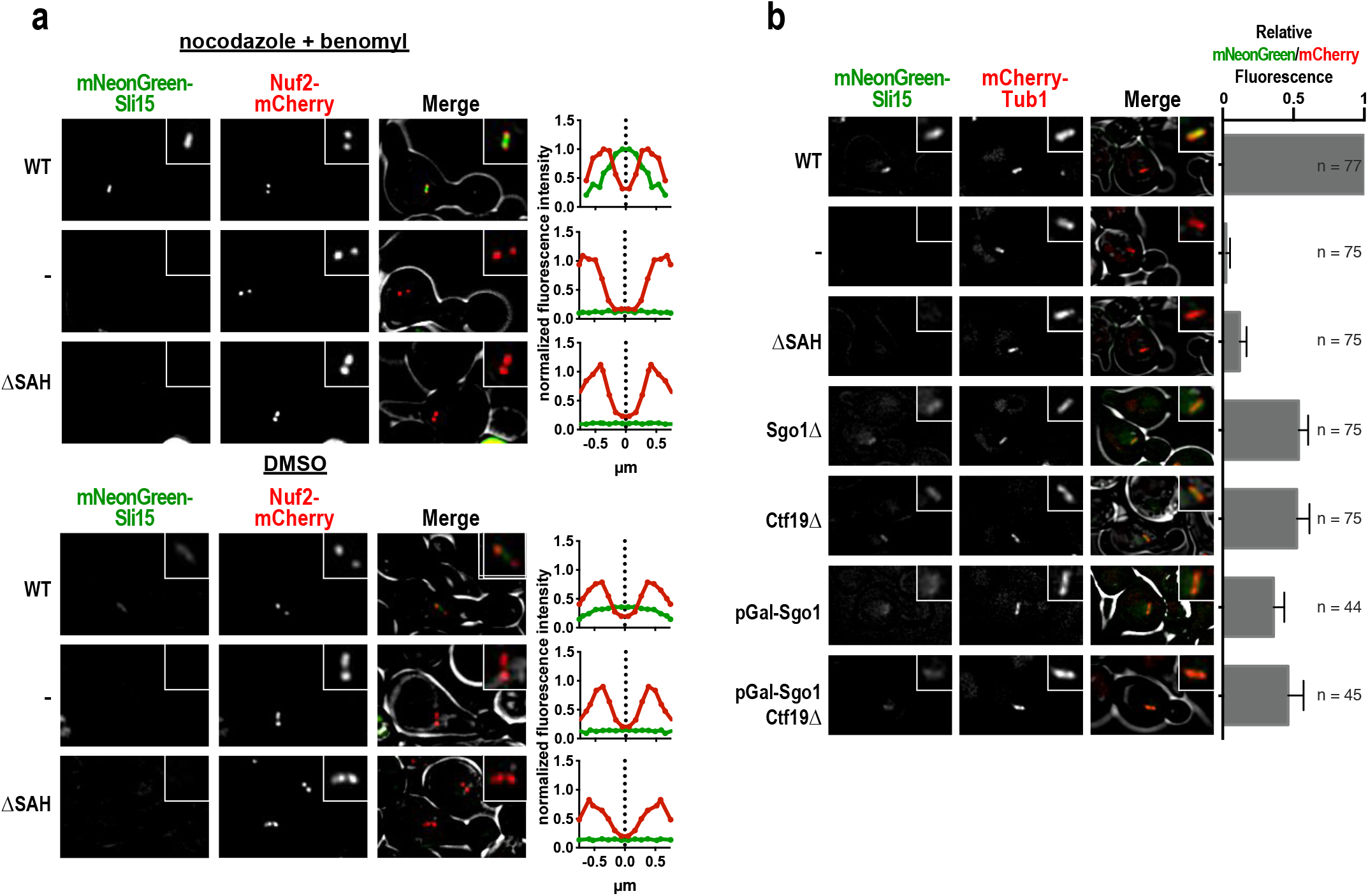
The SAH region of Sli15 contributes to its inner centromere and inner kinetochore localization. (**a**) Inner centromere localization of mNeonGreen-WT and mNeonGreen-ΔSAH was determined in preanaphase after 15 minutes of treatment with nocodazole (5 μM) and benomyl (34 μM) or DMSO. Nuf2-mCherry signal was used to determine the mitotic stage. Images of the same fluorophore are contrast adjusted the same. Insert box: 2.4 μm square. Enrichment of Sli15 constructs was analyzed by drawing a line along the spindle axis (Nuf2-mCherry signals). The intensities of mCherry and mNeonGreen signals of 20 cells were averaged and normalized to WT in nocodazole/benomyl. (**b**) Spindle localization of mNeonGreen-WT in preanaphase with the indicated depletions and deletions. Quantification was performed as in Figure 2c. The plot shows averages of at least three independent experiments. Inserts of representative images are 2.2 μm squares.

We next determined how the disruption of CPC localization in the Sli15-ΔSAH mutant compares to specific perturbation of CPC recruitment to the inner centromere or inner kinetochore. Deletion of either Sgo1 or Ctf19 each decreased CPC spindle localization by ~50% (Figure 5b). By comparison, Sli15-ΔSAH fluorescence decreased by ~90%. Simultaneous removal of Ctf19 and Sgo1 by deletion and depletion, respectively, still had more spindle fluorescence than Sli15-ΔSAH. This residual signal in the double deletion/depletion strain is potentially due to the microtubule binding activity of the SAH. This suggests that the CPC spindle localization signal observed in preanaphase results from a combination of inner centromere, inner kinetochore and microtubule binding, and that the SAH region is necessary for all three of these activities.

### Inner centromere localization is sufficient for error correction by the CPC

The lack of any measurable localization of the CPC in preanaphase for the SAH mutants presented us with an opportunity to individually restore targeting activities and determine which localizations are sufficient for CPC function at the outer kinetochore. We have already shown that microtubule targeting is not sufficient to rescue viability (Figure 3a), so we next determined if inner centromere, inner kinetochore, or outer kinetochore targeting could restore error correction. Sli15 constructs were fused to Sgo1 (inner centromere), Okp1 (inner kinetochore, COMA complex), or Dad3 (outer kinetochore, Dam1 complex) and expressed from the Sli15 promoter. All three fusion constructs distinctly change the localization pattern of Ipl1 from the wild-type localization pattern evenly distributed along the spindle. On average, the fusion constructs instead displayed two intensity peaks at different positions relative to the outer kinetochore marker Nuf2-mCherry (Figures 6a and S3a). To determine whether tethering of different Sli15 mutants to the inner centromere or inner kinetochore would rescue CPC function, the viability of these chimeras was tested in Sli15-depleted cells. Fusion of Sli15-12Q or Sli15-ΔSAH to Sgo1 led to a partial rescue of viability (Figure 6b), suggesting that recruitment of the CPC to the inner centromere is sufficient for biorientation. Expression of Sgo1-Sli15-ΔSAH or Sgo1-Sli15-12Q did not lead to a strong metaphase arrest (Figure 6c), indicating that the partial growth phenotype is likely due to low CPC activity as opposed to continuous kinetochore-microtubule attachment turnover. In contrast to the Sgo1 fusion, fusion of Sli15-ΔSAH to Okp1 or Dad3 did not rescue viability (Figures 6d and S3b). The Okp1 fusion also failed to increase Ndc80 phosphorylation in preanaphase, indicating that the observed lethality results from too little CPC activity (Figure S3c).

**Figure 6.**
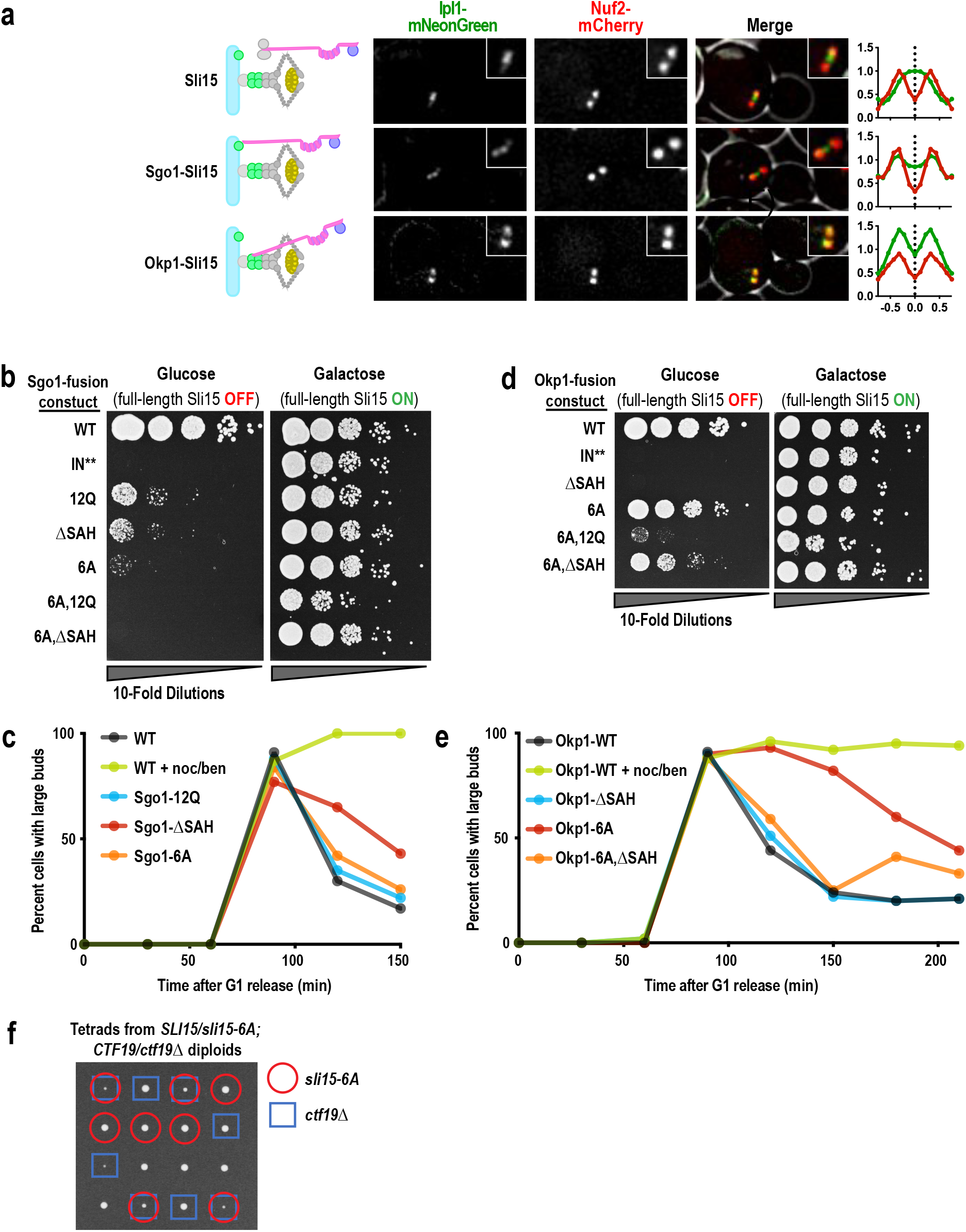
Tethering the CPC to the inner centromere or inner kinetochore and microtubules partially rescues deletion of the SAH. (**a**) Images for localization of CPC-fusions to Sgo1 or Okp1 in preanaphase cells. Cells were grown asynchronously for 2h in YPAD before imaging. Fluorescent signal was quantified as in Figure 5A. Values from 20 cells were averaged for each condition. All images with the same fluorophore were contrast adjusted the same. Insert box: 2.1 μm. (**b**, **c**) Serial dilutions of indicated Sli15 mutants fused to Sgo1 or Okp1, respectively. Endogenous Sli15 was under the control of a galactose inducible promoter. IN** denotes two point mutations in the IN box that prevent Ipl1 binding. (**d**, **e**) Budding index measurements of cells expressing the indicated Sli15 fusion constructs. Cells were arrested in G1 and endogenous Sli15 was depleted. The percentage of cells with large buds was determined every 30 min after release. Addition of nocodazole (5 μM) and benomyl (34 μM) to Okp1-Sli15 was used as a control for metaphase arrest. (**f**) Spores from SLI15/sli15-6A; CTF19/ctf19Δ diploid. Note that the combination of sli15-6A and ctf19Δ leads to a synthetic growth defect.

### Inner kinetochore and microtubule binding by the CPC synergize to establish chromosome biorientation

To determine if kinetochore or centromere targeting act synergistically with microtubule binding, we engineered Sli15-6A,ΔSAH double mutant fusion constructs. For the inner kinetochore (Okp1) targeting, this construct now partially rescued viability, demonstrating cooperativity between inner kinetochore and microtubule binding for CPC function (Figure 6d). Surprisingly, increased microtubule binding eliminated the rescue seen in the inner centromere-targeting Sgo1 fusion, suggesting incompatibility between the inner centromere and microtubule binding activities. These opposing effects of microtubule binding either complementing or conflicting with inner kinetochore versus inner centromere targeting were observed even in the absence of the ΔSAH mutation. Fusion of Sgo1 to the 6A mutation had a synthetic lethal phenotype that did not result in a cell cycle delay, indicating a lack of CPC activity. (Figure 6b, compare to Figure 3a). In contrast to Sgo1-Sli15-6A, cells expressing the Okp1-Sli15-6A construct have a robust mitotic delay, as would be expected for overactive CPC (Figure 6e). Sensitivity to the microtubule depolymerizing drug benomyl suggests that chromosome biorientation is partially compromised in this construct (Figure S3d). To test whether the delay was due to activation of the spindle assembly checkpoint (SAC), we combined the Okp1-Sli15-6A with a deletion of the checkpoint protein Mad2. Indeed, Mad2 deletion fully eliminated the mitotic delay. This suggests that the observed mitotic delay stems from over-activation of the checkpoint, potentially due to the continuous formation of unattached kinetochores caused by overactive kinetochore phosphorylation (Figure S3d). The opposing effects of microtubule binding on inner centromere vs. inner kinetochore localization suggest that the two recruitment pathways act independently. In agreement with this conclusion, combining Sgo1-Sli15-ΔSAH with Okp1-Sli15-6A,ΔSAH, had an additive effect on growth when compared to the single fusion constructs (Figure S3e).

To test for cooperativity between inner kinetochore targeting and microtubule binding independently of the Okp1-fusion, we determined if inner kinetochore targeting becomes more crucial for biorientation when microtubule binding is increased. Spores from heterozygous *SLI15/sli15-6A* and *CTF19/ctf19Δ* diploids demonstrated a clear synthetic negative genetic interaction between *sli15-6A* and deletion of the inner kinetochore protein Ctf19 (Figure 6f). Consistent with this, we previously found that a Sli15 mutant lacking the N-terminal CEN box and has increased spindle localization also shows a synthetic negative interaction with deletion of Ctf19 (Campbell and Desai, 2013).

As the combination of microtubule binding and inner kinetochore binding rescue SAH deletion, we tested whether these two targeting methods could rescue in trans. Simultaneous expression of constructs with microtubule targeting (Sli15-6A,ΔSAH) and inner kinetochore targeting (Okp1-Sli15-ΔSAH) failed to rescue growth, whereas a single construct with both targeting activities does rescue (Figure S3g). This indicates that microtubule and inner kinetochore binding of the CPC act synergistically on the same molecule (Figure S3g). Similarly, the negative effects of combining inner centromere targeting and microtubule targeting were not observed when the two constructs were expressed in trans (Figure S3f). The incompatibility between these two binding activities could therefore result from the inability of a single molecule to bind to both regions at the same time. Together, these data demonstrate that either localization to the inner centromere or to the inner kinetochore plus microtubules is sufficient to provide enough error correction to support cell viability.

## Discussion

In this study, we have used mutations in the SAH region of the INCENP/Sli15 subunit of the CPC to determine its contribution to chromosome segregation fidelity in budding yeast. These SAH mutants are lethal and result in extremely high rates of chromosome missegregation, demonstrating that the SAH is essential for chromosome biorientation. Accordingly, SAH disruption causes a substantial decrease in CPC substrate phosphorylation specifically at the kinetochore. In agreement with these results, SAH deletion leads to a decrease in kinetochore phosphorylation and mitotic checkpoint activity in human cell culture (Wheelock et al., 2017; Vader et al., 2007).

We find that the SAH region contributes to the localization of the CPC at the inner centromere, inner kinetochore, and microtubules during chromosome biorientation. A first indication for this conclusion came from the lack of any clear enrichment of CPC mutants harboring SAH mutations during this point of the cell cycle. By comparison, specific disruption of these known targeting mechanisms individually leads to a relatively mild decrease in localization signal. Localization of the CPC to the microtubules probably results from the strong overall net positive charge of the SAH region. Recruitment to the inner kinetochore likely occurs through a direct interaction between CTF19 and a region of Sli15 adjacent the SAH (Fischböck-Halwachs et al., 2019). The requirement of the SAH to localize to the inner centromere is difficult to explain, as it would not be predicted to disrupt CPC recruitment via the Sli15 CEN box. However, a decrease in chromatin association in HeLa cells and frog egg extracts was also observed for SAH-deleted INCENP, suggesting that this property of the SAH is conserved (Wheelock et al., 2017).

We did not detect any CPC localization in the SAH mutants in preanaphase even after the disruption of microtubule binding at the kinetochore to promote inner centromere localization. This allowed us to use these mutants to then restore the targeting to specific regions and determine if any of the known CPC interactions were sufficient for activity. Although microtubule binding is the best-characterized property of SAH domains across species, restoration of this activity through compensatory mutations was not sufficient to rescue biorientation. Intriguingly, microtubule binding does lead to outer kinetochore phosphorylation, even in cells with bioriented attachments. This result strongly suggests a tension-independent targeting of kinase activity to attached kinetochores via microtubule binding. Recently published results for a Borealin-based microtubule binding activity suggest that a similar mechanism occurs in human cells (Trivedi et al., 2019). We find that outer kinetochore phosphorylation by microtubule targeting alone did not lead to SAC activation, implying that low levels of phosphorylation across all attached kinetochores does not lead to detachment. The role of microtubule binding may therefore be to increase the phosphorylation of syntelically-attached versus unattached kinetochores, both of which are not under tension.

The best-characterized and most clearly conserved preanaphase localization of the CPC is at the inner centromere. Accordingly, this is the only localization that was partially sufficient to restore viability in the SAH mutants independently of the other localization activities. However, we previously showed that the inner centromere localization of the CPC is not essential for accurate chromosome segregation, suggesting the existence of another pathway that is also sufficient for chromosome segregation in budding yeast (Campbell and Desai, 2013). Recent publications in human cells have also demonstrated that the inner centromere localization of the CPC is not essential for outer kinetochore phosphorylation (Hadders et al., 2020; Broad et al., 2020; Liang et al., 2020). Although neither inner kinetochore targeting nor microtubule targeting of the CPC were alone sufficient to rescue viability, the combination of the two partially suppressed the lethal phenotype to a similar extent as the inner centromere targeting. We therefore conclude that the combination of inner kinetochore and microtubule binding activities of the SAH region form a “kinetochore-intrinsic” pathway for chromosome biorientation.

Intriguingly, directing the CPC towards the inner centromere or the inner kinetochore has opposing affects when combined with increased microtubule binding. Directing the complex to inner centromere binding either through deletion of inner kinetochore protein Ctf19 or by direct tethering to the inner centromeric protein Sgo1 has a synthetic negative effect in combination with the Sli15-6A mutant that promotes microtubule binding. By contrast, promotion of inner kinetochore binding by direct tethering to the COMA complex member Okp1 results in synthetic rescue when microtubule binding is increased. Furthermore, we have previously demonstrated that an N-terminal deletion of Sli15 that also promotes microtubule binding fully rescues the growth of Sgo1 deletion mutants (Campbell and Desai, 2013). These data argue against a model wherein all three binding activities act simultaneously on the same CPC molecule. Instead, our data support two independent pathways of CPC recruitment (Figure 7).

**Figure 7.**
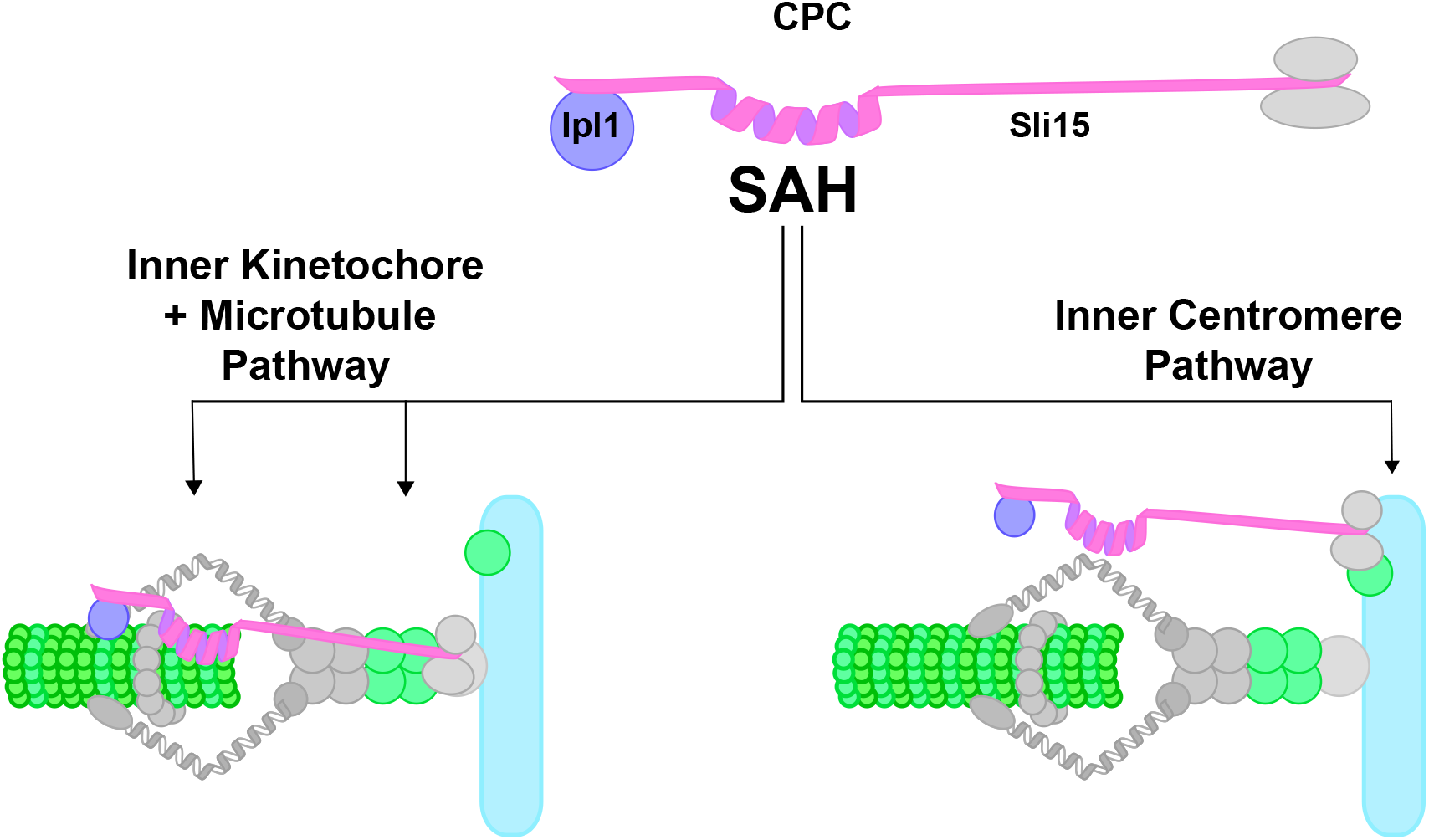
Two parallel pathways establish chromosome biorientation. Model for how CPC binding activities contribute to chromosome biorientation. CPC binding sites at the microtubule, inner kinetochore, and inner centromere are indicated in green. The inner centromere targeting forms one pathway, while the inner kinetochore and microtubule binding act cooperatively to form a second pathway. At syntelic attachments, all three binding activities contribute to creating very high levels of kinetochore phosphorylation, selectively destabilizing such attachments.

It remains to be determined if these two pathways completely overlap in their activities or if they have some functional specificity. The CPC is involved in both kinetochore formation and the regulation of kinetochore-microtubule attachments and has different substrates for each of these functions (Akiyoshi et al., 2013; Rago et al., 2015; Kim and Yu, 2015). The two pathways could differ in their contributions to these two activities (Bonner et al., 2019). Alternatively, the two pathways could preferentially regulate different types of misattachments.

## Materials and methods

### Yeast strains and media

All yeast strains used were grown in yeast extract/peptone containing 40 μg/ml adenine-HCl (YPA) and 2% glucose (YPAD) or 1% raffinose and 1% galactose (YPAGR). Unless otherwise stated, cultures were grown at 30°C. Galactose inducible promoters, fluorescent tags, epitope tags and gene deletions were introduced into the genomic loci as previously described (Longtine et al., 1998). Sli15 constructs were cloned into plasmids via Gibson assembly (Gibson et al., 2009). Plasmids were integrated at the URA3 or TRP1 loci by digesting the plasmids with BstBI or BsgI, respectively. All integrations and plasmids were checked by sanger sequencing.

### Yeast transformation

PCR products or digested plasmids were transformed into cells pelleted from 50 mL exponentially growing cultures. Cells were washed once with TE (10 mM Tris, pH 7.5 and 1 mM) EDTA and once with LiAc/TE (100 mM LiAc, 10 mM Tris, pH 7.5 and 1 mM EDTA). Cells were resuspended in LiAc/TE and 100 μL were added to 10 μL plasmid or PCR product, 10 μL single stranded sheared salmon sperm DNA and 750 μL PEG/LiAc/TE (40% PEG 4000, 100 mM LiAc, 10 mM Tris and 1 mM EDTA, pH 7.5). After 30 min at 30°C 80 μL DMSO were added and incubated at 42° C for 15 min Then cells were pelleted, resuspended in SOS (0.3 YPAD /YPAGR and 6 mM CaCl_2_), plated on selective plates and incubated for two days at 30°C. For resistance markers G418 or hygromycin B cells were grown overnight at 30°C on YPAD / YPAGR and replica-plated the next day on the respective selective plate.

### Yeast synchronization and galactose inducible depletion

To deplete proteins under the control of a *Gal10-1* promoter, cells were synchronized for 45 min in YPAGR containing α-factor (10 μg/mL). Media was exchanged with YPAD containing α-factor and grown for an additional 2 h and 15 min Cells were then released into the cell cycle by washing twice with YPAD and adding Pronase E (Merck). Nocodazole (VWR) was used at a concentration of 5 μM and benomyl (Sigma-Aldrich) was used at 34 μM. Rapamycin (Santa Cruz Biotechnology) was used at a concentration of 1 μM.

### Protein extraction and Western blotting

For protein extraction, saturated overnight cultures were diluted into 5 mL YPAGR to an OD of 0.25. After synchronization and depletion of Sli15 and/or Cdc20, cells were released into the cell cycle and, unless otherwise stated, harvested 60 min after release. Proteins were extracted by pelleting the cells, resuspending in 100 μL trichloroacetic acid (5%) and incubating them for 10 min at room temperature. Cells were washed once with 1 mL ddH_2_O and resuspended in 100 μL lysis buffer (50 mM Tris, pH 7.4, 50 mM dithiotreitol, 1 mM EDTA, cOmplete EDTA-free protease inhibitor cocktail (Roche) and Phosstop (Roche). Glass beads were added and the cells were vortexed for 30 min at 4°C. 33 μL 4x sample buffer were added and incubated at 95°C for 3 min The samples were stored at −20° C. To measure Sli15 expression, cells were harvested, pelleted and resuspended in 100 μL NaOH (0.2 M), pelleted and resuspended in 1x sample buffer before incubating them at 95° C for 3 min and stored at −20°C. For immunoblots, the following antibodies were used: mouse anti-Pgk1 monoclonal 22C5D8 (Thermo Fisher Scientific), rat anti-HA-clone 3F10 (Roche), mouse anti-Myc monoclonal 4A6 (EMD Millipore), rabbit anti-p-Histone H3 (Ser 10) polyclonal (Santa Cruz Biotechnology), mouse anti-Maltose Binding Protein monoclonal antibody IgG2a (New England Biolabs), mouse anti-Histone H3 [Trimethyl Lys9] 6F12-H4 (Novus Biologicals), rabbit anti-Ndc80 phospho-S37 polyclonal. Phosphospecific antibodies were used with 40 μM unphosphorylated competitor peptide. Membranes were then probed with the corresponding secondary antibodies: anti-rat IgG-HRP-linked (Cell Sgnaling Technology), anti-rabbit IgG-HRP-linked (Cell Sgnaling Technology) or anti-mouse IgG-HRP-linked (Cell Sgnaling Technology). Immunoblots were quantified using ImageJ (National Institutes of Health).

### Microscopy

Unless otherwise stated, cells were synchronized as described above and imaged 60 min or 90 min after release. Cells were pelleted and washed twice with 1 mL sterile ddH_2_O, resuspended in 100 μL sterile ddH_2_O and 2 μL of cell suspension were spotted onto 1% agarose pads supplemented with complete synthetic media and 2% glucose. A coverslip was placed on top and sealed with VALAP (1:1:1 mixture by weigth of paraffin (Merk), lanolin (Alfa Aesar) and Vaseline (Ferd. Eimermacher). Images were collected on a DeltaVision Ultra Epifluorescence Microscope system (Cytiva) at 23°C and a PlanApo N 60/1.42 Oil objective and a sCMOS sensor, 2040 x 2040 pixels, 6.5 μm pixel size camera. 12 z-sections with step size of 0.5 μm were taken, except for the chromosome missegregation assay where 24 z-sections were taken. Images were deconvolved using softWoRx software (Life Sciences Software). Quantifications and subsequent normalizations were performed from images obtained on the same day and representative images were contrast adjusted identically using ImageJ. For spindle localization in preanaphase and anaphase cells, ImageJ was used to measure the intensity distribution of a perpendicular line (5 pixels wide) to the spindle center. The obtained intensities were fitted to a gaussian curve and subsequently integrated as previously described (Fink et al., 2017). For this analysis non-deconvolved images were used. For line-scan analysis intensity distributions of a line (5 pixels wide) along the spindle axis were background subtracted and the average of 20 cells were calculated.

### Recombinant protein expression and purification

Sli15 microtubule binding domain constructs were designed with an MBP (maltose binding protein) tag at the N terminus and a 6X Histidine tag at the C terminus. All proteins were expressed in *Escherichia coli* Rosetta pLysS. Overnight cultures were each inoculated in 2L 2XTY medium containing 50 μg/mL kanamycin and grown at 37°C to OD600 of 1. Protein expression was induced at 18°C by 0.3 mM IPTG for 15 hours. Harvested cells were resuspended in lysis buffer (50 mM Tris pH 7.4, 300 mM KCl, 10% Glycerol, 1 mM DTT) containing one cOmplete Mini EDTA free protease inhibitor cocktail tablet and 0.2 mM PMSF and lysed by sonication. Lysates were centrifuged at 14000 rpm at 4°C and the supernatants clarified with a 0.2 μm membrane syringe filter (VWR International) before proceeding with a two-step purification protocol. Clarified lysates were passed on preequilibrated Histrap HP columns (GE Healthcare). The columns were washed with wash buffer (lysis buffer with 40 mM Imidazole) and the bound proteins were eluted in incremental fractions with 100, 150 and 200 mM imidazole. Pooled eluted fractions were concentrated with Ultracel-30 (Millipore) and injected on pre-equilibrated Superdex HiLoad S200 16/60 or 26/600 (GE Healthcare) at 4°C with lysis buffer. Protein fractions were collected and frozen in liquid nitrogen and stored at −80°C.

### Microtubule Pelleting assay

Porcine brain tubulin (Cytoskeleton, Inc) reconstituted with tubulin buffer (80 mM PIPES pH 6.9, 1 mM MgCl_2_, 0.5 mM EGTA, supplemented with GTP) was thawed and spun in an ultracentrifuge (Beckman Coulter OptimaMax) at 70k rpm for 5 min, 4°C to remove insoluble protein. To polymerize tubulin, 1 mM GTP was added, 1 and 10 μM taxol were sequentially added and incubated at 37°C for 10 min each. After incubation at 37°C for another 15 min, the microtubules were added to a 40% glycerol cushion containing tubulin buffer, GTP and taxol and spun in an ultracentrifuge at 37°C for 20 min. The pellet was rinsed with mQ and resuspended in tubulin buffer with 20 μM taxol to yield 10 μM polymerized microtubules.

For pelleting assays, recombinant MBP fusion proteins purified from *E. coli* were thawed from −80°C and first spun at 70k rpm for 30 min at 4°C in an ultracentrifuge. Supernatants were carefully separated and diluted in buffer containing 50 mM Tris pH 7.4, 150 mM KCl, 10% glycerol and 1 mM DTT. 25 nM proteins were added to 1 μM taxol-stabilized microtubules in 50μl reactions and incubated for 10 min at 25°C. All reactions contained 0.2 mg/mL BSA. Pellets and supernatants were separated by ultracentrifugation as above at 25°C and heated with SDS sample buffer at 95°C for 5 min.

### Tetrad dissection

Diploid strains were grown on YPAD plates overnight at 30°C, streaked onto sporulation plates (16 mg/mL sodium acetate and 16 mg/mL agar) and incubated for 2 days at 23°C. Cells were then treated with zymolase (Zymo Research) for 15 min at 30°C and dissected onto YPAD plates. For genotyping, spores were replica plated onto appropriate selective plates.

### Statistical analysis

Samples that were compared to each other were obtained in the same experiment. Statistical significance between samples was assessed via an unpaired two-sided Student’s *t* test using Prism 6 (Graphpad).

### Computer code and code availability

The python script used in Figures 5a, 6a and S3 to determine the localization of the CPC relative to Nuf2 was developed for this study. Briefly, from deconvolved images, a line (5 pixels wide) along the Nuf2-mCherry signals was drawn and fluorescence signal intensities along this line were obtained for mCherry and mNeonGreen. After background subtraction, a signal intensity threshold of the mCherry signal defined the length of the spindle. The obtained position and corresponding intensity values for mCherry and mNeonGreen were averaged for all spindles analyzed from the same sample. Normalization to WT and data plotting was performed in Excel (Microsoft) and Prism 6 (Graphpad), respectively. The code is available by the authors upon request.

## Acknowledgements

The authors thank the Campbell and Dammermann laboratories as well as Franz Klein for helpful discussions and comments on the manuscript. We thank Peter Schlögelhofer, Franz Klein and Gustav Ammerer for gifts of yeast strains and plasmids. We acknowledge the service of the MPL Central Biooptics-Light Microscopy, a member of VLSI.

This work was supported by Vienna Science and Technology Fund (WWTF) grant VRG14-001 and Austrian Science Fund (FWF) grants Y944-B28 and W1238-B20 to C.S.C.

## Author contributions

P.Y. performed the experiments in Figure 2a. C.M. performed experiments in Figure 1f and Figure S1e. K.S. contributed to experiments, cloning, strain production and establishment of the Microtubule pelleting assay. T.M. performed the rest of the experiments. P.Y., C.M., K.S., T.M. and C.S.C. conducted data analysis. C.S.C. and T.M. wrote the manuscript. C.S.C. supervised the project.

---

**Figure S1.**
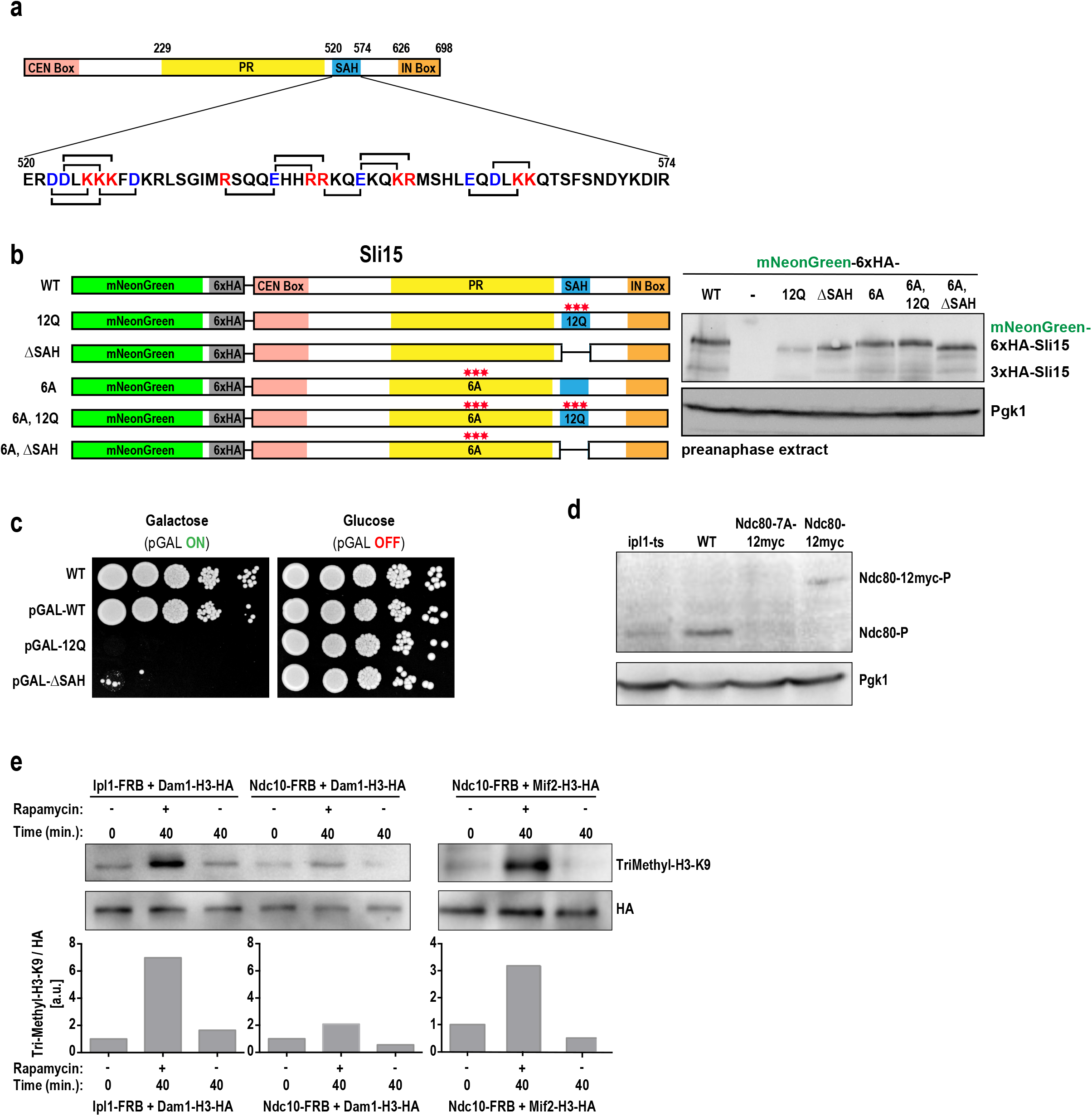
Related to Figure 1. (**a**) Schematic of Sli15 and amino acid sequence of the SAH region. Amino acids of opposite charge in the configuration i±3 and i±4 are highlighted in blue (acidic) or red (basic). (**b**) Immunoblot of Sli15 construct expression levels in preanaphase when endogenous Sli15 was depleted. Expression levels were tested using an antibody against the HA epitope. Sli15 constructs were labelled with mNeonGreen and 6xHA tags whereas endogenous Sli15 was labelled with 3xHA. Pgk1 is shown as loading control. (**c**) Serial dilution of 12Q and ΔSAH overexpression. Overexpression of the Sli15 constructs under the control of a galactose inducible promoter was induced by growing the cells on galactose containing plates. (**d**) Immunoblot showing the specificity of the Ndc80-S37 phospho-specific antibody. This antibody was generated for this study based on published literature (Akiyoshi et al., 2009; Fink et al., 2017). Extracts were prepared from ipl1-321, Ndc80 WT, Ndc80-12myc and Ndc80-12myc bearing alanine mutations at Ipl1 phosphorylation sites. Pgk1 is shown as a loading control. (**e**) Immunoblots to determine the spatial resolution of the M-track assay. Ndc10, a centromeric protein, is proximal to Mif2 located at the inner kinetochore, but not proximal to the outer kinetochore protein Dam1. Signal intensities were normalized to the 0 timepoint without rapamycin addition.

**Figure S2.**
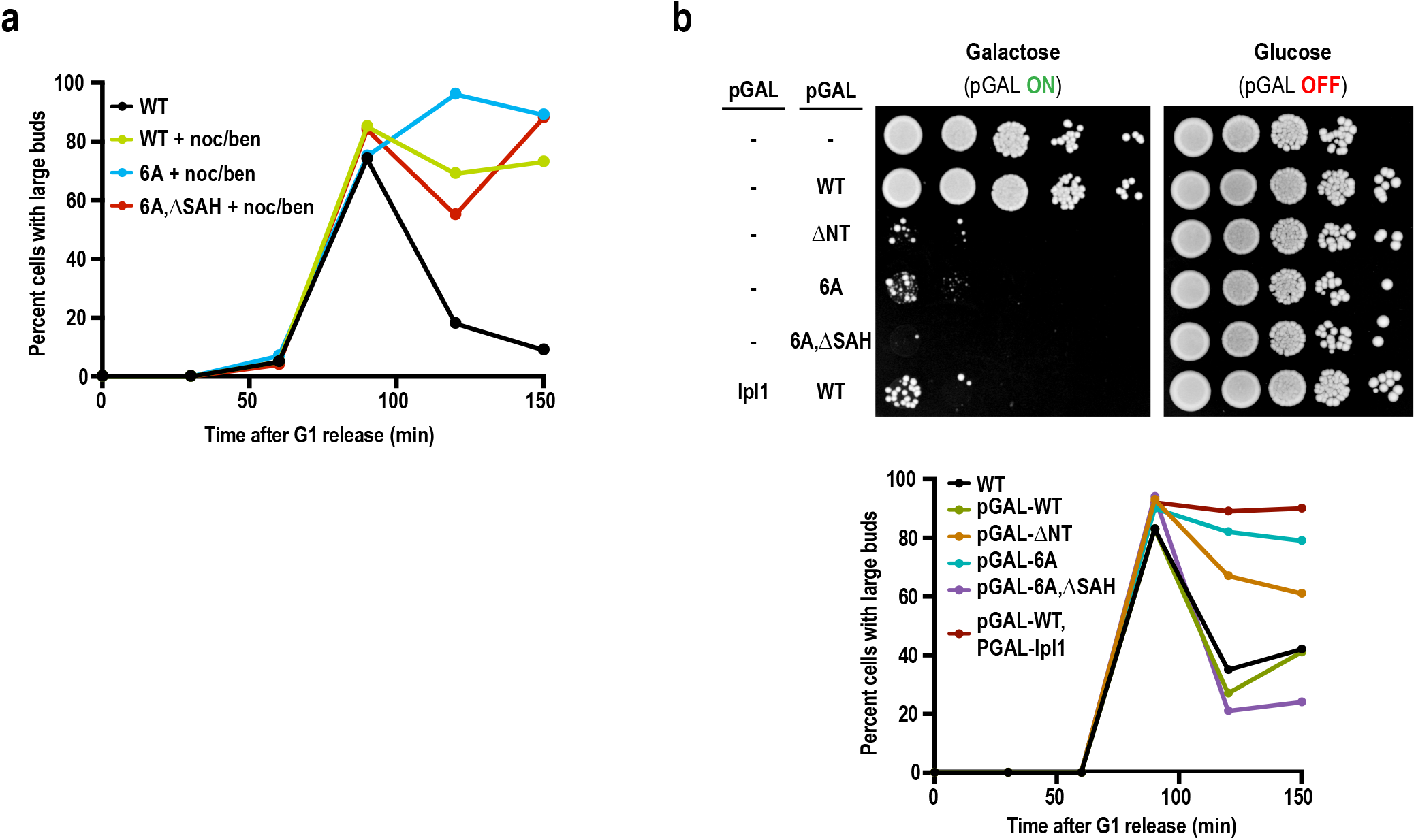
Related to Figure 4. (**a**) Budding index analysis of WT, Sli15-6A and Sli15-6A, ΔSAH. Cells were synchronized in G1 and depleted of the endogenous Sli15 before release. Nocodazole (5μM) and benomyl (34 μM) were added 45 min after release. WT without nocodazole/benomyl treatment served as a control for cell cycle exit. (**b**) Serial dilution and budding index measurements of the indicated strains. Sli15 constructs and Ipl1 were overexpressed from a galactose inducible promoter similar as described in Figure S1d. ΔNT is a truncated version of Sli15 with deletion of aa 2-228 (Campbell and Desai, 2013)

**Figure S3.**
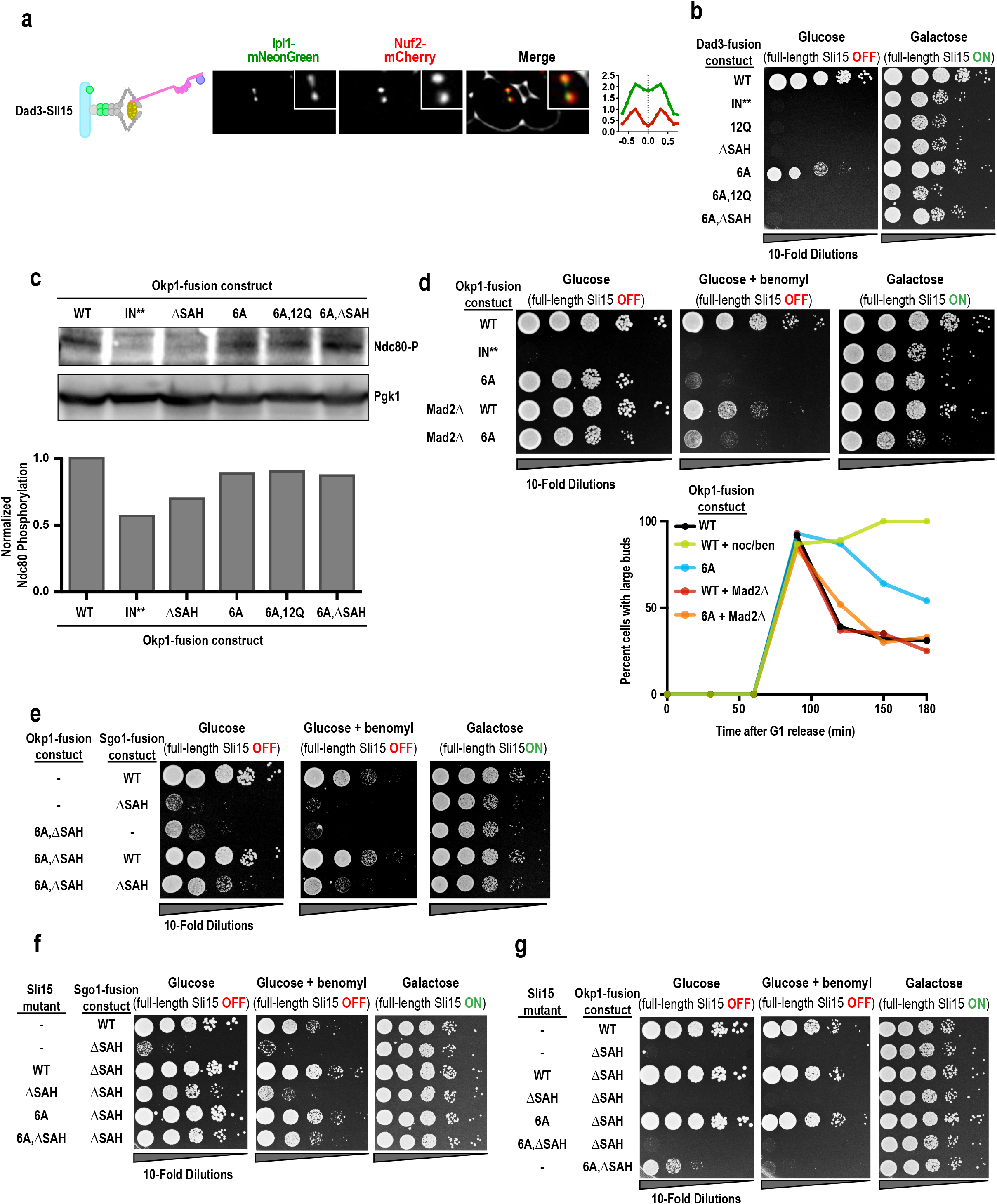
Related to Figure 6. (**a**) Localization of the CPC with Sli15 fused to Dad3. Line-scan analysis for mCherry and mNeonGreen signal distribution was performed as in Figure 5a. (**b**) Serial dilution analysis for indicated Sli15 constructs fused to Dad3. Endogenous Sli15 was under the control of a galactose inducible promoter and depleted on the glucose containing agar plate (left). (**c**) Immunoblot and quantification of Ndc80-S37 phosphorylation with strains containing the indicated Sli15 constructs fused to Okp1. Extracts were prepared from cells in preanaphase, previously synchronized in G1 while endogenous Sli15 was depleted. Pgk1 is shown as a loading control. (**d**) Serial dilution analysis and budding index measurements of indicated Okp1-Sli15 fusion constructs together with Mad2 deletion. Cells were synchronized in G1 and depleted of the endogenous Sli15 before release. As a control for metaphase arrest, Nocodazole (5μM) and benomyl (34 μM) were added to wild-type cells 45 min after release. (**e**) Serial dilution analysis of the combination of Okp1-Sli15 and Sgo1-Sli15 constructs. Cells were spotted on glucose, glucose + benomyl (10 μg/ml) and galactose containing agar plates. (**f**) Serial dilution analysis of the combination of Sgo1-Sli15-ΔSAH and diffusible Sli15 constructs. (**g**) Serial dilution analysis of the combination of Okp1-Sli15-ΔSAH and diffusible Sli15 constructs.

**Table S1.** Related to Figure 2. Table showing the quantification of mCherry and mNeonGreen signals from the experiment in Figure 2b. Values are normalized to WT. Mean and SD of three independent experiments are shown. Note that the mCherry signal intensities only change marginally between different Sli15 constructs.

| | WT | - | 12Q | ΔSAH | 6A | 6A, 12Q | 6A, ΔSAH |
| --- | --- | --- | --- | --- | --- | --- | --- |
| **mCherry-Tub1** | 1 | 0.78±0.02 | 0.88±0.01 | 0.92±0.02 | 0.95±0.01 | 0.88±0.01 | 0.78±0.1 |
| **mNeonGreen-Sli15** | 1 | 0.009±0.006 | 0.3±0.1 | 0.3±0.2 | 4±1 | 3.3±0.9 | 2.5±0.1 |

**Table S2.**
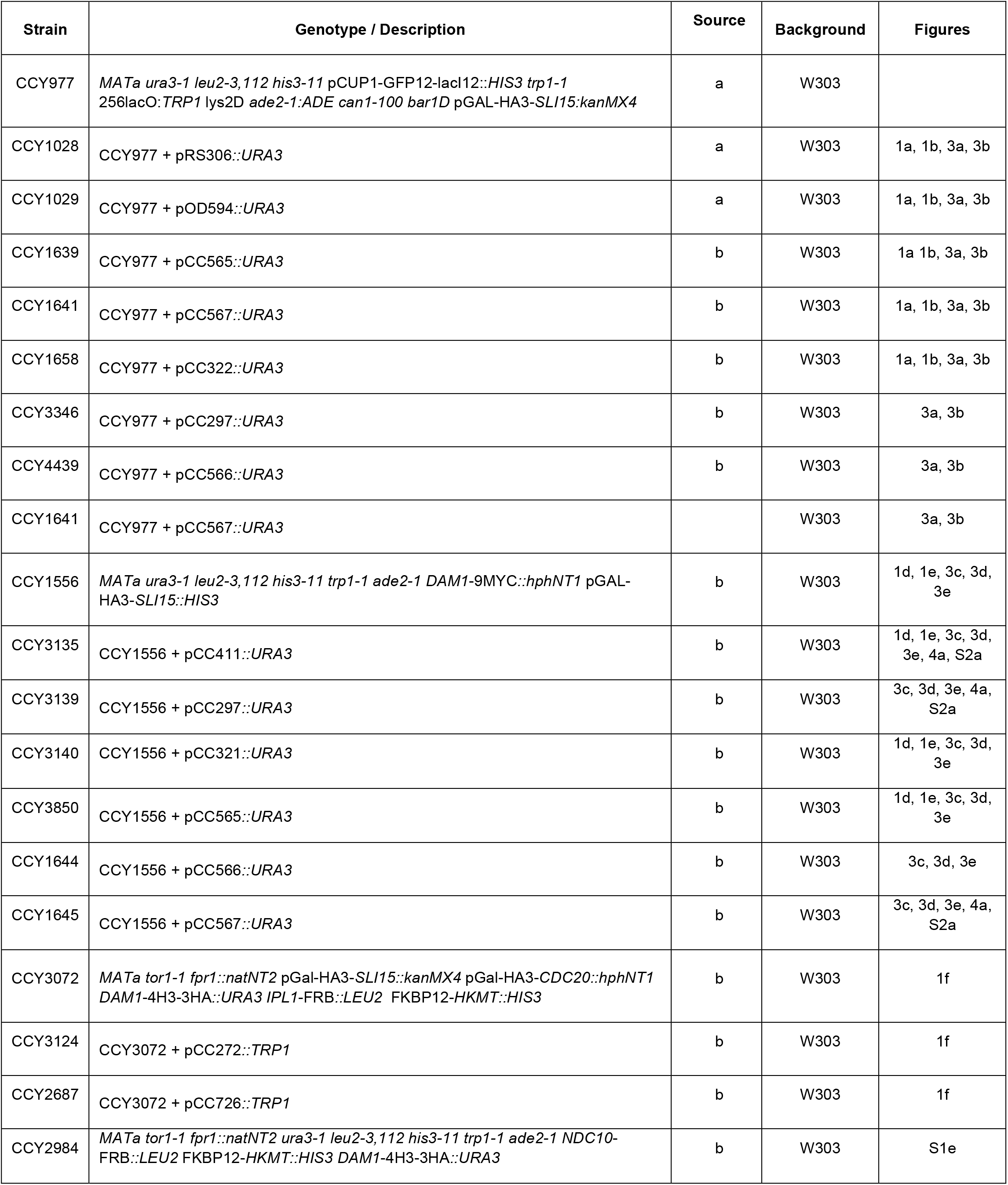
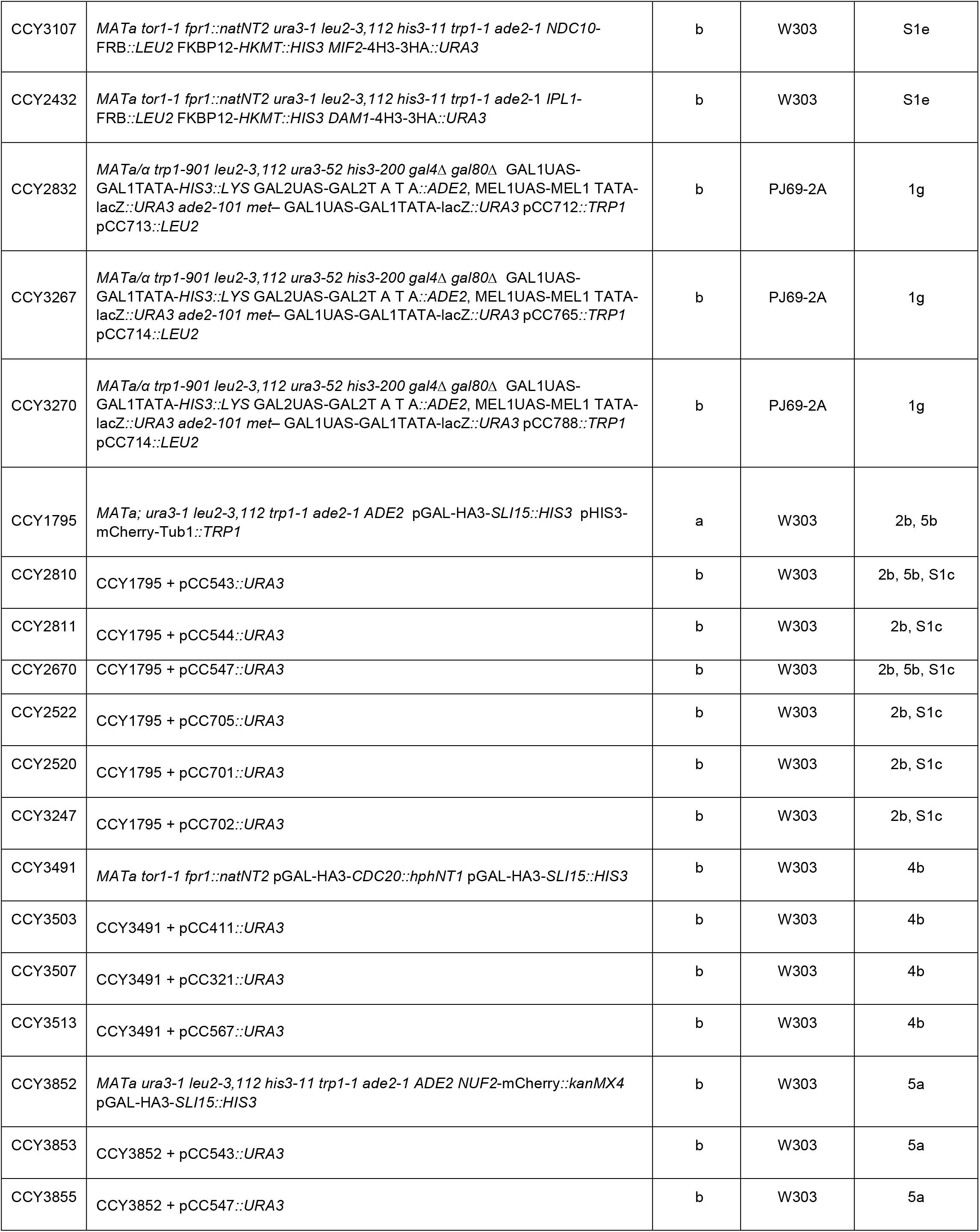
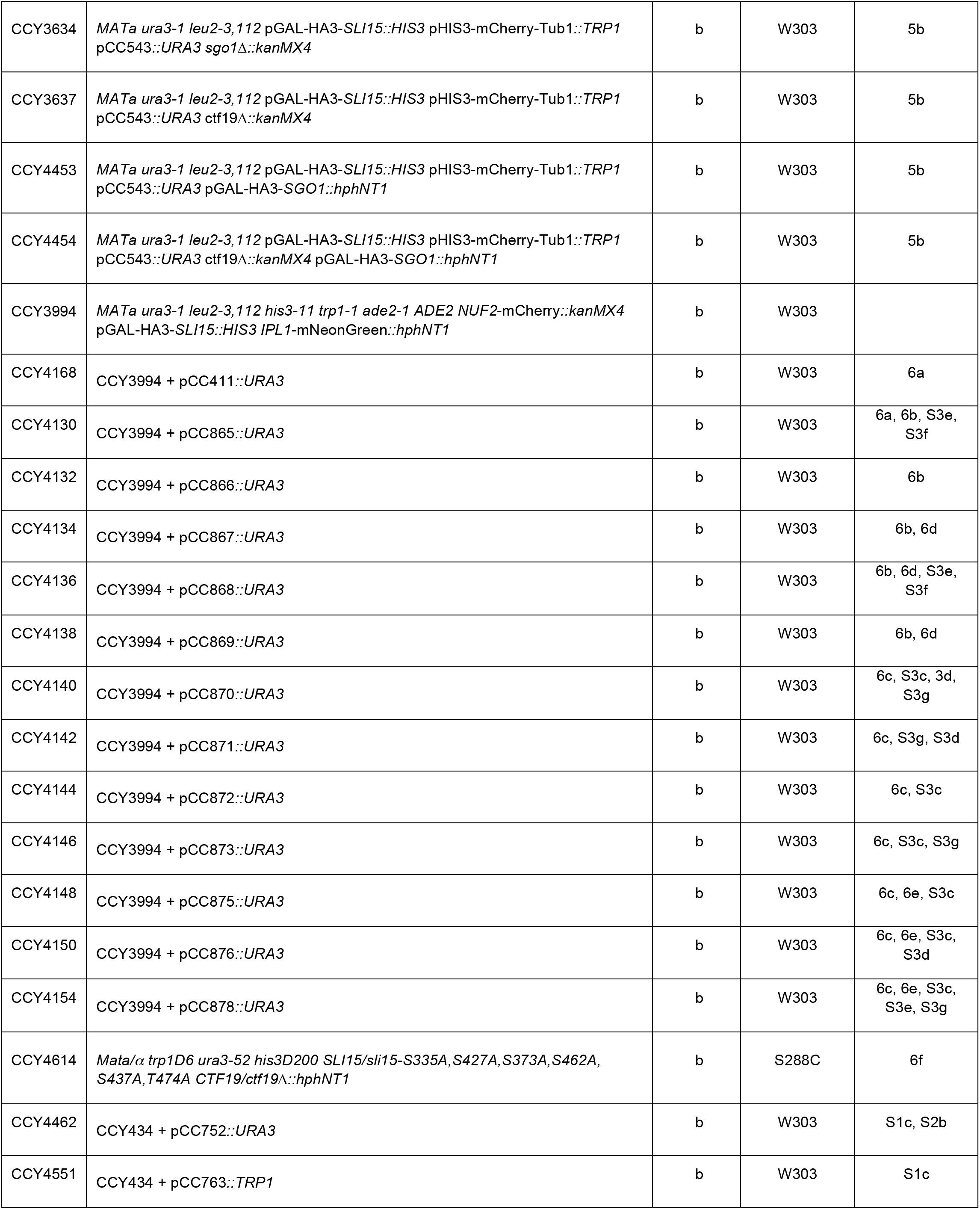
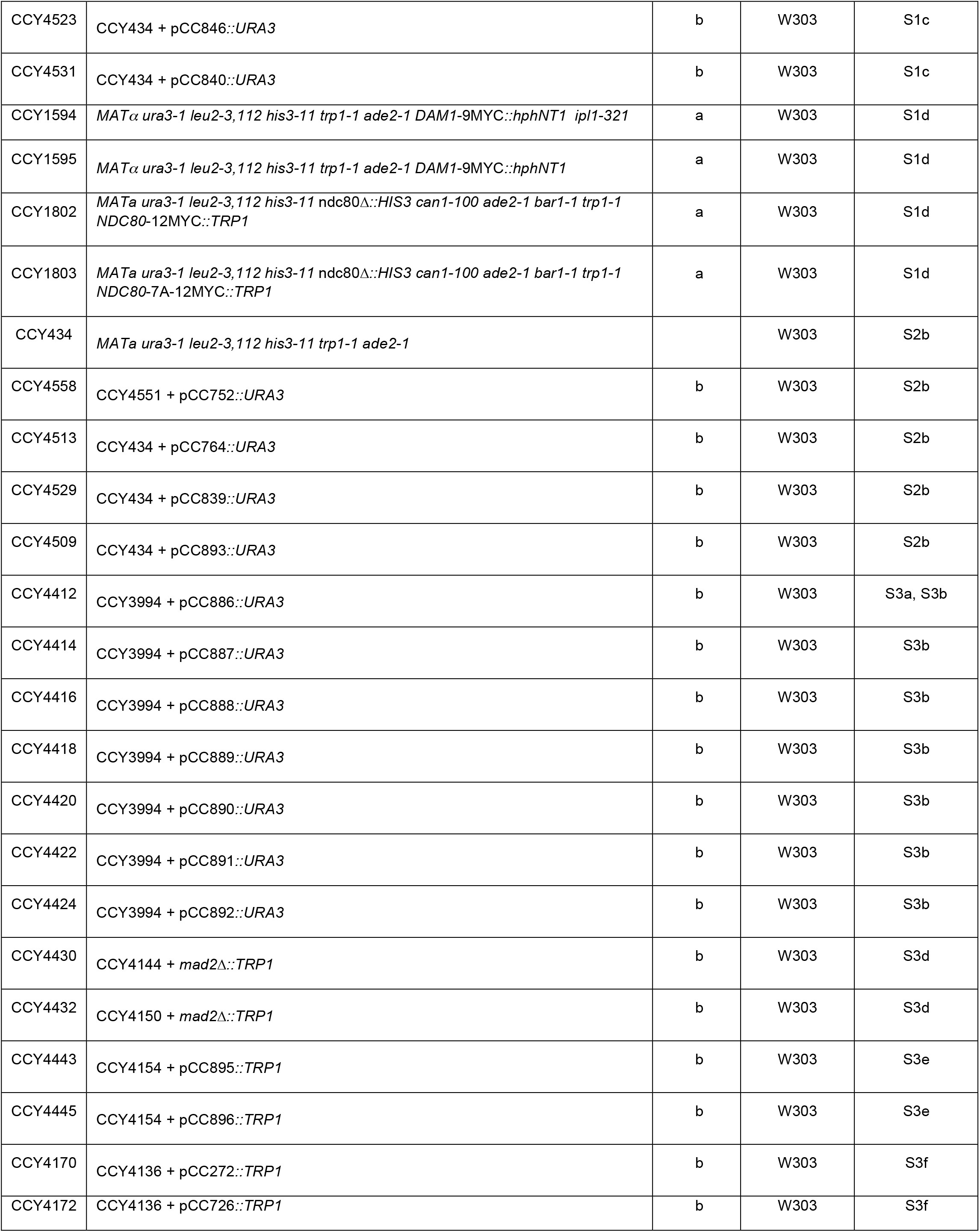
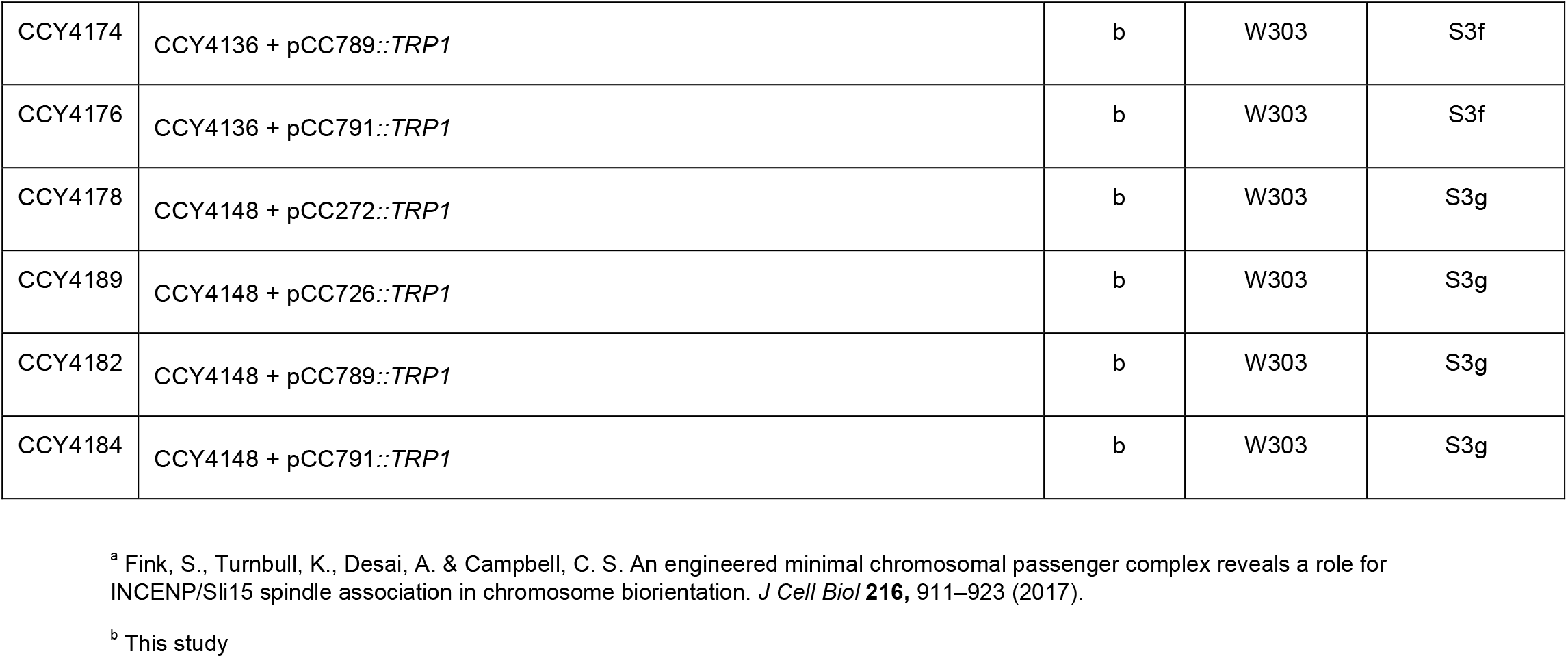
Yeast strains used in this study

**Table S3.**
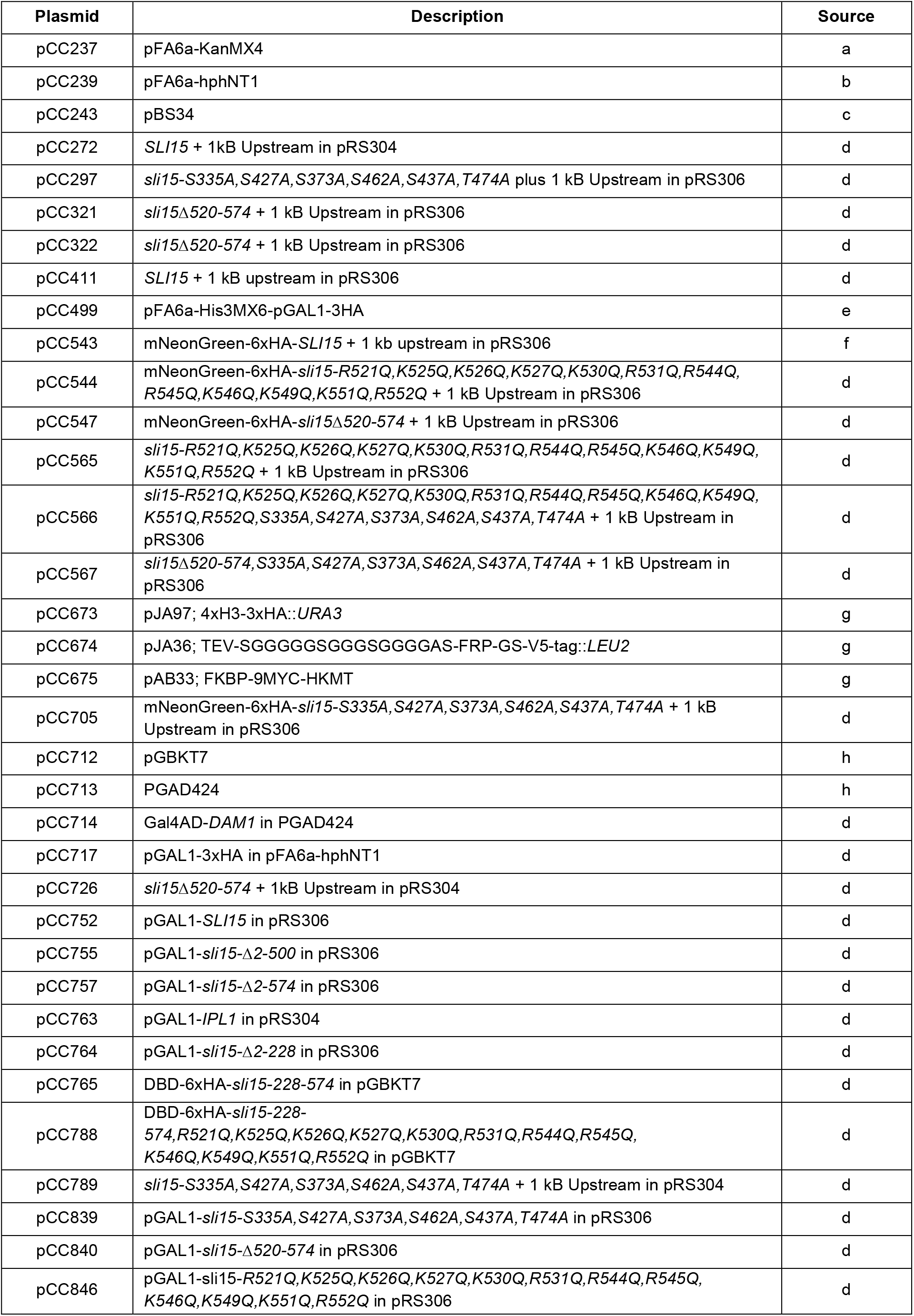
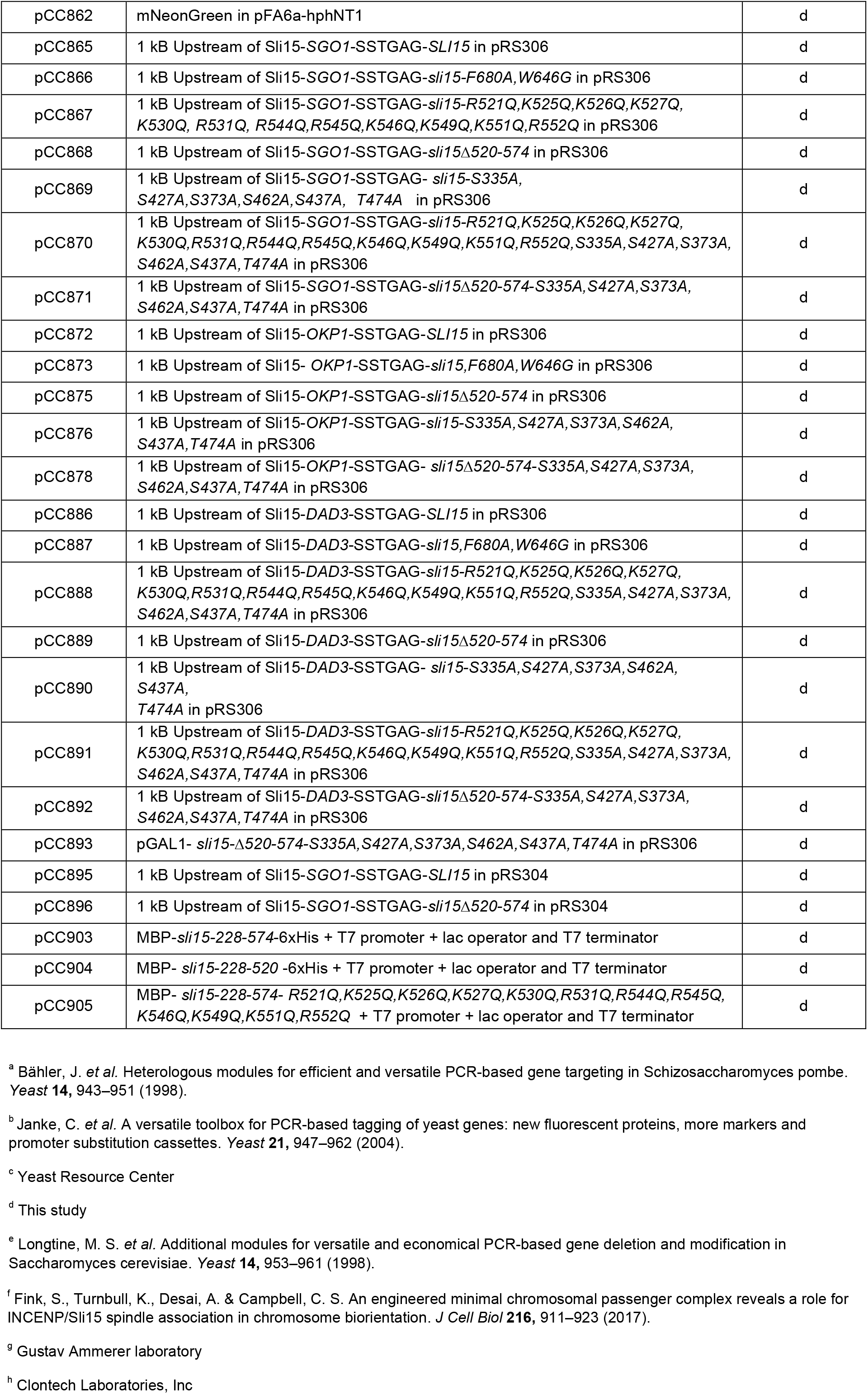
Plasmids used in this study

## Abbreviations

CPC: chromosomal passenger complex
PR: phosphoregulated
SAH: single alpha helix
MTB: Microtubule Binding
FRB: FKBP12 Rapamycin binding
HKMT: histone lysine methyltransferase
DBD: DNA Binding Domain
AD: Activation Domain

